# Robust and Adaptive Non-Parametric Tests for Detecting General Distributional Shifts in Gene Expression

**DOI:** 10.1101/2025.03.06.641952

**Authors:** Fanding Zhou, Alan J. Aw, Dan D. Erdmann-Pham, Jonathan Fischer, Yun S. Song

## Abstract

Differential expression analysis is crucial in genomics, yet existing methods primarily focus on detecting mean shifts. Variance shifts in gene expression are well-documented in studies of cellular signaling pathways, and more recently they have characterized aging, thus motivating the need for flexible detection approaches that include tests of expression variance changes. In this work, we present QRscore (Quantile Rank Score), a general method for detecting distributional shifts in gene expression by extending the Mann-Whitney test into a flexible family of rank-based tests. Here, we focus on implementing QRscore to detect shifts in mean and variance in gene expression, using weights designed from negative binomial (NB) and zero-inflated negative binomial (ZINB) models to combine the strengths of parametric and non-parametric approaches. We show through simulations that QRscore not only achieves high statistical power while controlling the false discovery rate (FDR), but also outperforms existing methods in detecting variance shifts and mean shifts. Applying QRscore to bulk RNA-seq data from the Genotype-Tissue Expression (GTEx) project, we identified numerous differentially dispersed genes and differentially expressed genes across 33 tissues. Notably, many genes have significant variance shifts but non-significant mean shifts. QRscore augments the genome bioinformatics toolkit by offering a powerful and flexible approach for differential expression analysis. QRscore is available in R, at https://github.com/songlab-cal/QRscore.

## 1 Introduction

In both bulk and single-cell genomics, differential expression analysis is a routine procedure for identifying genes whose expressions differ under varying conditions or treatments. To detect changes in average expression, widely used methods in bulk RNA-seq differential analysis treat the dispersion of gene expression as a nuisance parameter, implementing techniques to shrink and stabilize the estimation of variance. The widespread use of these approaches can result in missed signals of gene expression variability that capture important biological mechanisms. In humans and *C. elegans*, variability indices are associated with gene regulation[1, 2] and cell development [3], potentially explaining many biological phenomena, including adaptation to changing environments, disease susceptibility, and incomplete penetrance [4, 5]. Recent studies also demonstrate that variability indices provide insights into aging [6, 7, 8], therefore underscoring the crucial need to incorporate changes in variability into tests of differential gene expression.

To detect expression variability, previous studies on microarray datasets [9, 10] applied traditional hypothesis testing methods, such as the F-test and Bartlett’s test. These tests are based on normal distribution assumptions of two samples and are designed to identify differences in standard deviation. To improve robustness, one approach is to analyze different measures for variability other than the standard deviation. Such measures include the median absolute deviation [10] and the coefficient of variation [2]. Another approach to ensure the robustness of statistical tests is to employ methods that are more robust to deviations from normality for two samples, such as Levene’s test and permutation tests [10]. The method diffVar [11] implements a modified version of Levene’s test that utilizes an empirical Bayes estimator instead of variance estimation for *t*-statistics to obtain a more powerful test. However, this method maintains its parametric nature, making it susceptible to reduced performance due to deviations from the underlying assumptions. Additionally, non-parametric methods like the Kolmogorov-Smirnov (KS) test are not commonly used in the analysis of RNA-seq data because of their limited statistical power, especially when working with small sample sizes.

Because bulk RNA-seq data often show overdispersion, particularly in gene expression where the variance for most genes exceeds the mean, they are commonly modeled using the negative binomial (NB) or zero-inflated negative binomial (ZINB) distribution. One method for detecting differences in dispersion applies generalized additive models for location, scale, and shape (GAMLSS) [12], allowing the addition of regressions for dispersion parameters in the NB distribution into the main regression model, and targeting changes in dispersion parameters through likelihood ratio tests [4]. Another method, MDseq [13], utilizes a reparametrization of the ZINB model to test the mean and dispersion separately from generalized linear models (GLMs). DiffDist [14] uses a hierarchical Bayesian model based on the NB distribution, which can provide tests for differential dispersion and distribution for RNA-seq data.

Although these current methods are based on negative binomial distributions and are widely applied in scenarios of small sample sizes, recent research has shown that parametric methods are prone to inflated False Discovery Rate (FDR) in both studies with small sample sizes [15] and population-level RNA-seq studies with large sample sizes [16]. Additionally, they are highly sensitive to preprocessing procedures [17] and are not robust to model misspecification [18].

Here, we introduce QRscore (Quantile Rank Score), a flexible rank-based method for conducting robust and powerful two-sample tests. QRscore generalizes the Mann-Whitney test and capitalizes on the adaptability of the chosen test statistic to yield substantial statistical power across various scenarios, including the detection of shifts in mean and variance. QRscore offers various approaches for designing test statistics that are effective in detecting shifts in mean and dispersion. These include a versatile naive approach applicable to various cases beyond count data, as well as methods designed specifically for RNA-seq count data with either a low occurrence of zeros or a higher proportion of inflated zeros. Through simulations that extensively compare QRscore with existing differential variability tests and differential centrality tests, we find that QRscore controls false discovery rates across diverse model specifications, while simultaneously obtaining high statistical power. We use QRscore to detect both differentially dispersed genes (DDGs) with variance shifts and differentially expressed genes (DEGs) with mean shifts across multiple tissues in the Genotype-Tissue Expression (GTEx) project [19], showing *differential variability* characterizes functionally distinct sets of genes than *differential centrality*. Our method is further expanded to include *K*-sample tests, making it suitable for multi-group scenarios. Our research contributes to the field of bioinformatics tools for genomics, and our methods can be adapted to other applications that require flexible non-parametric tests.

## 2 Results

### 2.1 Overview of QRscore

QRscore is a non-parametric, rank-based hypothesis testing method for detecting differential mean or differential dispersion in RNA-seq data. This method leverages the strengths of non-parametric tests to obtain robust false positive control, while benefiting from parametric modeling to increase detection power when the data closely follow a parametric model (Fig. 1).

**Figure 1.**
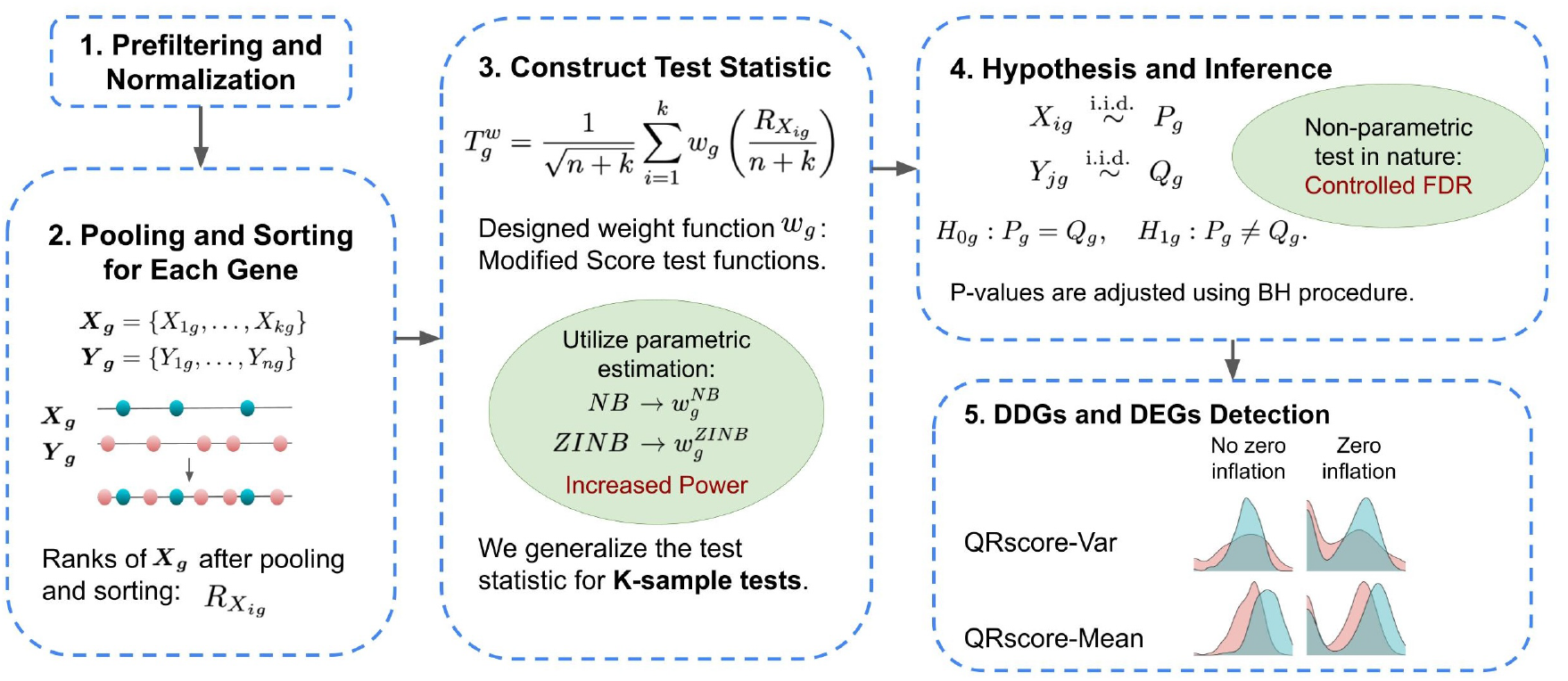
Overview of QRscore: (1) Prefiltering and Normalization - This initial data processing stage ensures quality and comparability of data; (2) Pooling and Sorting - For gene *g*, expression levels *X*_*g*_ = {*X*_1*g*_, …, *X*_*kg*_} and *Y*_*g*_ = {*Y*_1*g*_, …, *Y*_*ng*_} are combined into a single ordered list. The rank of *X*_*ig*_, denoted as 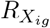,is determined after pooling and sorting the expression values of gene *g*; (3) Constructing the Test Statistic - A test statistic *T* is calculated using a weighted function based on modified score test functions, which enhances test power through parametric estimation and incorporates a generalized *K*-sample test; (4) Hypothesis and Inference - This stage involves evaluating the null hypothesis for gene *g, H*_0*g*_ : *P*_*g*_ = *Q*_*g*_, where *P*_*g*_ and *Q*_*g*_ represent the distributions of gene expression samples *X*_*g*_ and *Y*_*g*_, respectively. *p*-values are computed and adjusted using the Benjamini-Hochberg procedure to control the false discovery rate; (5) Identification of DDGs and DEGs - Genes with significant expression level differences between the conditions are identified, as represented in the QRscore-Var and QRscore-Mean diagrams. Details of QRscore-Var and QRscore-Mean are provided in Appendix A.

Our null hypothesis posits that the two samples are drawn from the same distribution. Specifically, given a gene *g*, let *X*_*ig*_ and *Y*_*jg*_ denote the observed read counts for individual *i* and *j* in the first and second conditions, respectively, where *i* = 1, …, *k* and *j* = 1, …, *n*. With samples 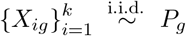 and 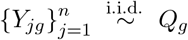,the null and alternative hypotheses for gene *g* are:

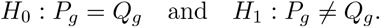

Our rank-based test statistic, 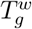,is computed as the normalized sum of weighted ranks of the *X*_*ig*_ observations, using the formula:

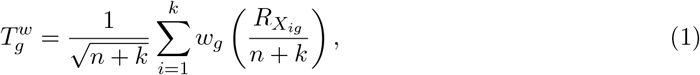

where 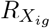 denotes the rank of *X*_*ig*_ after pooling and sorting observations from both groups. Here, *w*_*g*_ is a weight function chosen to maximize the test power against user-specific alternatives. Its dependence on *g* indicates that it can be chosen differently across different genes.

While the space of all possible weight functions *w*_*g*_ induces a family of test statistics of the form Eq. (1), particular choices of *w*_*g*_ lead to well-known tests. For instance, setting *w*_*g*_(*x*) = *x* transforms Eq. (1) into one equivalent to the Mann-Whitney test statistic. Intuitively, Mann-Whitney assigns higher weights to larger ranks, thereby enhancing the power to detect differences in means. On the other hand, if we choose a quadratic weight function 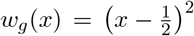 that places greater weights on the extremes and less in the middle, the test is well-powered for assessing changes in variance (this weighting scheme corresponds to Mood’s test of scale [20, Section 4]).

Focusing on differential expression analysis applications, we design weight functions *w*_*g*_ by composing the score function (the gradient of the log-likelihood with respect to the parameters) with the inverse cumulative distribution function (Methods, Section 4.1). We derive these score functions selectively from either a NB or a ZINB distribution, with QRscore-Var targeting DDGs and QRscore-Mean targeting DEGs. The rationale behind this design is to overcome the lower power of traditional rank-based models; our approach significantly boosts power under correct model specification and achieves efficiency comparable to the likelihood ratio test.

Despite the tailored weight design, our method remains distribution-free under the null hypothesis *H*_0_, which assumes identical distributions (*P*_*g*_ = *Q*_*g*_). This is achieved because the rank-based inputs are uniformly distributed under the null, regardless of the underlying data distribution. This distribution-free property provides a significant advantage: it inherently reduces the impact of model misspecification and helps maintain strict control over false discoveries.

Moreover, our method also extends to support *K*-sample tests, broadening its applicability to multiple sample scenarios. Overall, QRscore is a versatile and powerful approach for detecting differential mean or dispersion in RNA-seq data, combining the robustness of non-parametric tests with the enhanced detection power of special weight design. The details of the test statistic construction and its statistical inference are described in Methods.

### 2.2 Simulation results

In this section, we perform extensive simulations to evaluate existing methods for detecting genes with significant changes in dispersion or mean across groups. To benchmark 2-sample tests, we use mean and dispersion parameters estimated from real RNA-seq count data of different age groups from whole blood tissue in the GTEx project as the baseline. We then introduce varying levels of fold changes in dispersion or mean for randomly selected DDGs. In the differential dispersion analysis, each simulated dataset applies uniform fold changes—either increases or decreases—across the selected DDGs, with the magnitude of these changes varying between datasets. We also simulate datasets with different proportions of genes with outliers or inflated zeros to assess the impact of these factors on method performance. Additionally, we evaluate the effect of sample size by altering the number of individuals in each age group, thus testing the methods under conditions that closely mimic actual experimental settings. Finally, we also compare our method in pairwise 2-sample tests with 3-sample tests for DDG analysis by simulating 3 samples with different fold changes and sample sizes. Simulation details are in Section 4.2.

Key simulation results are qualitatively summarized in Table 1. The hypothesis tests can be categorized into two groups: those that rely on specific parametric distributions and those that are non-parametric. For DDG analysis, parametric methods include GAMLSS, MDSeq, diffVar, and Levene’s test, while Mood and QRscore-Var are distribution-free. Similarly, for DEG analysis, methods like DESeq2, edgeR, and limma-voom rely on parametric models, whereas Mann-Whitney and QRscore-Mean are distribution-free. To analyze the sensitivity and robustness of these methods, metrics such as Area Under the Precision-Recall Curve (AUPRC), Power, and FDR under different simulation settings are computed. Non-parametric methods generally showed greater robustness to outliers and zeros. Notably, QRscore-Var and QRscore-Mean achieved the highest detection power for identifying DDGs and DEGs across various scenarios. In contrast, parametric methods were more susceptible to the negative effects of outliers and inflated zeros, often leading to an inflated FDR. Detailed results are provided in the following subsections.

**Table 1:** Summary table for simulation results.

| Method | Assumption | Targeted direction | Efficient in NB model | Robust to Outliers | Robust to Zeros |
| --- | --- | --- | --- | --- | --- |
| QRscore | Non-parametric | Mean/Variance | Yes | Yes | Yes |
| GAMLSS | NB | Variance | Yes | No | No |
| MDSeq | ZINB | Mean/Variance | No | No | Yes |
| Mood | Non-parametric | Variance | No | Yes | Yes |
| diffVar | Gaussian | Variance | Yes | Yes | No |
| Levene | Gaussian | Variance | Yes | Yes | No |
| DESeq2 | NB | Mean | Yes | No | Yes |
| edgeR | NB | Mean | Yes | No | Yes |
| limma-voom | Gaussian | Mean | Yes | Yes | No |
| Mann-Whitney | Non-parametric | Mean | Yes | Yes | Yes |

#### 2.2.1 QRscore is a solid contender in differential dispersion and differential expression analysis under correctly specified NB models

In simulations based on negative binomial distributions, QRscore-Var exhibits strong performance, slightly behind GAMLSS (Setting 1 in Table 2) and achieving comparably high detection power with FDR controlled (Fig. 2, “No Zero Inflation or Outliers”). This close competition highlights QRscore-Var’s effectiveness in scenarios with correctly specified NB models. QRscore-Var also demonstrates higher power than naive methods like Mood’s test and Levene’s test (Supplementary Fig. S1A). In smaller sample size settings (Setting 2-3 in Table 2), QRscore-Var consistently ranks as the second-best method after GAMLSS, reinforcing its reliability and showing notable improvements as sample sizes increase.

**Table 2:**
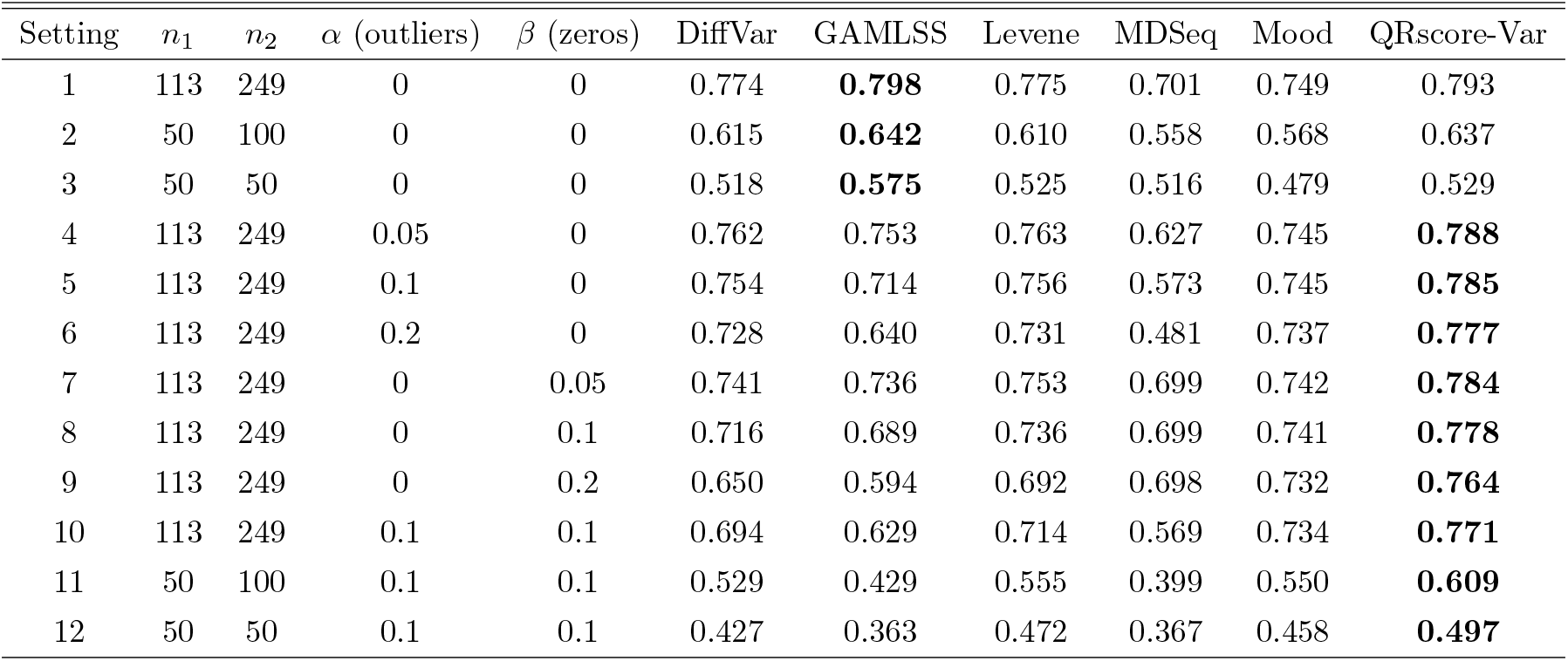
Average Area Under the Precision-Recall Curve (AUPRC) across all fold changes all seeds for each simulation setting. The dispersions of Group 1 are greater than those of Group 2 for true DDGs. For each setting, the greatest value is shown in bold. Sample size for Group 1: *n*_1_, sample size for Group 2: *n*_2_, proportion of genes with random outliers: *α*, proportion of genes with random zeros: *β*. Sample sizes *n*_1_ = 113 and *n*_2_ = 249 are chosen from GTEx whole blood tissue data, corresponding to 40-49 and 60-69 age groups respectively. When perturbations are introduced, QRscore outperforms the others, and it ranks second when model assumptions are perfectly met.

| Setting | $n_1$ | $n_2$ | $\alpha$ (outliers) | $\beta$ (zeros) | DiffVar | GAMLSS | Levene | MDSeq | Mood | QRscore-Var |
| --- | --- | --- | --- | --- | --- | --- | --- | --- | --- | --- |
| 1 | 113 | 249 | 0 | 0 | 0.774 | **0.798** | 0.775 | 0.701 | 0.749 | 0.793 |
| 2 | 50 | 100 | 0 | 0 | 0.615 | **0.642** | 0.610 | 0.558 | 0.568 | 0.637 |
| 3 | 50 | 50 | 0 | 0 | 0.518 | **0.575** | 0.525 | 0.516 | 0.479 | 0.529 |
| 4 | 113 | 249 | 0.05 | 0 | 0.762 | 0.753 | 0.763 | 0.627 | 0.745 | **0.788** |
| 5 | 113 | 249 | 0.1 | 0 | 0.754 | 0.714 | 0.756 | 0.573 | 0.745 | **0.785** |
| 6 | 113 | 249 | 0.2 | 0 | 0.728 | 0.640 | 0.731 | 0.481 | 0.737 | **0.777** |
| 7 | 113 | 249 | 0 | 0.05 | 0.741 | 0.736 | 0.753 | 0.699 | 0.742 | **0.784** |
| 8 | 113 | 249 | 0 | 0.1 | 0.716 | 0.689 | 0.736 | 0.699 | 0.741 | **0.778** |
| 9 | 113 | 249 | 0 | 0.2 | 0.650 | 0.594 | 0.692 | 0.698 | 0.732 | **0.764** |
| 10 | 113 | 249 | 0.1 | 0.1 | 0.694 | 0.629 | 0.714 | 0.569 | 0.734 | **0.771** |
| 11 | 50 | 100 | 0.1 | 0.1 | 0.529 | 0.429 | 0.555 | 0.399 | 0.550 | **0.609** |
| 12 | 50 | 50 | 0.1 | 0.1 | 0.427 | 0.363 | 0.472 | 0.367 | 0.458 | **0.497** |

**Figure 2.**
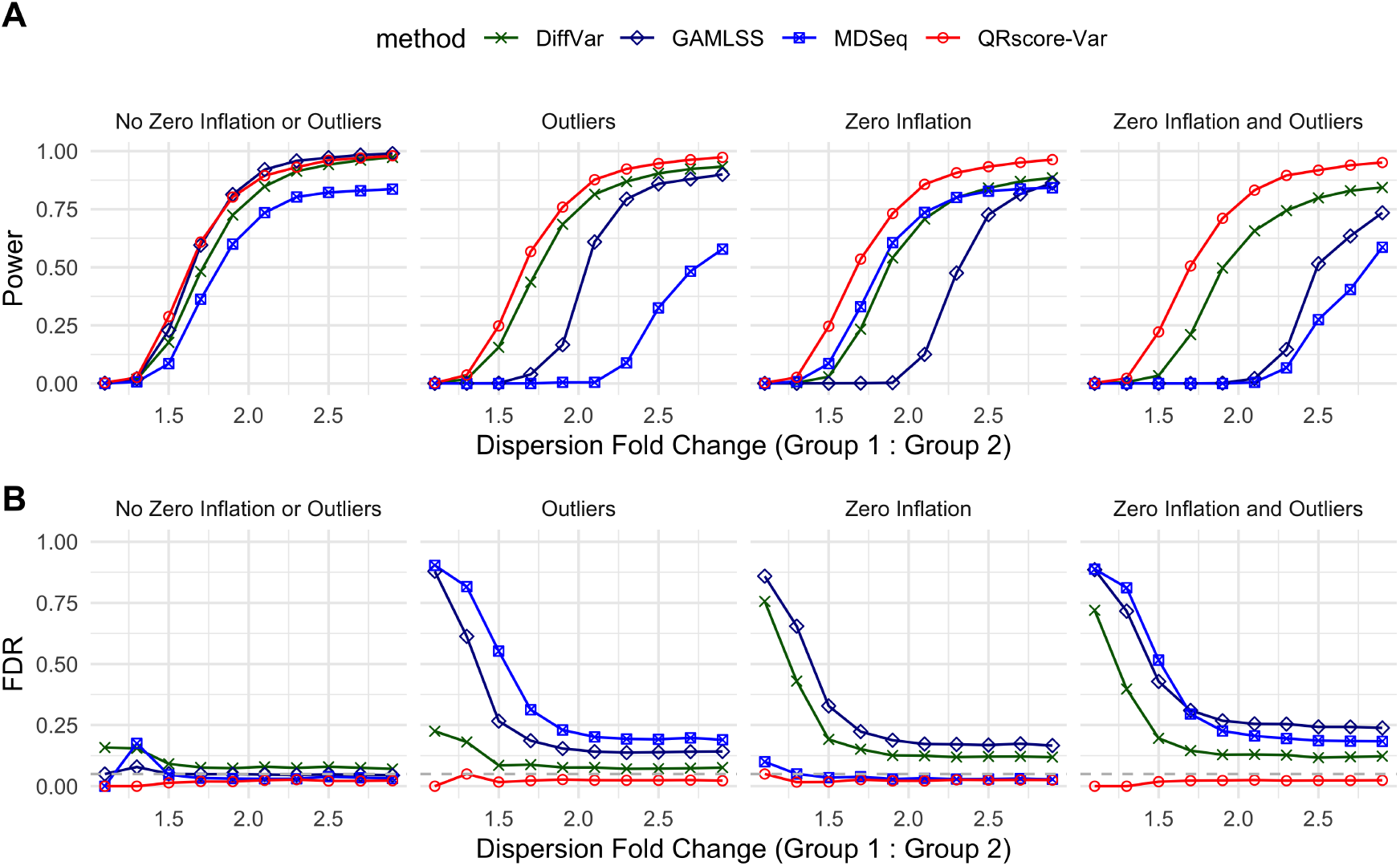
Simulation results for QRscore-Var benchmarked against computational biology methods for detecting DDGs. The dispersions of Group 1 are greater than those of Group 2 for true DDGs.**(A)** Line plots of power when true false discovery proportions are less than 0.05 across different dispersion fold changes. (**B**) Line plots of FDR when adjusted *p*-values are less than 0.05 across different dispersion fold changes. In **A** and **B**, we present the results for sample sizes *n*_1_ = 113 and *n*_2_ = 249. We present four different settings in the plot: one simulated from a pure NB model without random zeros or outliers; one with 10% of genes having outliers; one with 10% of genes having zero inflation; and one with both, where 10% of genes have outliers and 10% have zero inflation.

Likewise, as shown in Supplementary Table S2, QRscore-Mean shows performance comparable to DESeq2 and edgeR in differential mean analysis, with a marginal difference in AUPRC of about 0.5%. This slight difference in power and FDR is further illustrated in Setting 1 of Supplementary Fig. S2.

Furthermore, QRscore maintains a controlled Type I error rate, even in smaller sample scenarios, as shown in simulations based on pure NB distributions without introducing any fold change signals (Supplementary Fig. S3). This suggests that the need for slower exact testing procedures, such as resampling methods, is reduced in our study.

#### 2.2.2 QRscore-Var leads in robustness and effectiveness in gene expression variability analysis with outliers and excess zeros

Next, we simulate data with random outliers (Setting 4-6 in Table 2), where QRscore-Var emerges as the best performing method based on average AUPRC. In contrast, methods like GAMLSS and MDSeq show much lower AUPRC scores, even with a small proportion of outliers, as their reliance on the negative binomial assumption makes them vulnerable to model misspecification. For instance, in Setting 4 with 5% random outliers, the average AUPRC score for GAMLSS drops by 5.6% (from 0.798 to 0.753) compared to its performance in Setting 1, while QRscore-Var experiences only a 0.6% drop (from 0.793 to 0.788). As the proportion of outliers increases to 20%, GAMLSS’s score declines by 19.8% (to 0.640), while our method remains stable with only a 2.0% drop (to 0.777). Similarly, the parametric method MDSeq shows significant performance degradation in settings with outliers. This is further evidenced in scenarios with 10% outliers, where both GAMLSS and MDSeq experience an inflated FDR, as shown in Fig. 2B (“Outliers”).

In simulations with random zeros (Setting 7-9 in Table 2), QRscore-Var again demonstrates superior performance. While QRscore-Var handles excess zeros effectively, GAMLSS and diffVar falter under zero-inflated conditions. The poor performance of the latter two methods is highlighted by their inflated FDR and diminished power when controlling for the actual false discovery proportions in the scenarios with 10% random zeros (Fig. 2, “Zero Inflation”). Furthermore, QRscore-Var outperforms MDSeq across all fold changes, achieving higher power despite MDSeq’s ability to handle technical zeros effectively. This consistent high detection power and effective FDR control underscore QRscore-Var’s robustness in the zero-inflated scenarios.

In the context of decreased sample sizes with outliers and zeros present (Setting 11-12 in Table 2), QRscore-Var consistently achieves the highest average AUPRC regardless of the sample sizes. Additionally, while the previous results focus on scenarios where Group 1 has higher variance than Group 2 in true DDGs, we also conduct simulations where Group 2 has higher variance to evaluate performance under different variance change directions. The results showed that QRscore-Var maintained its top ranking compared to other methods when there are zeros or outliers present, further demonstrating its robustness (Supplementary Table S1, Supplementary Fig. S4, Supplementary Fig. S5).

#### 2.2.3 QRscore-Mean leads in robustness and effectiveness in differential gene expression analysis with outliers and excess zeros

QRscore-Mean shows the strongest performance in differential gene expression analysis under high proportions of random outliers (Setting 6 in Supplementary Table S2). While edgeR and DESeq2 show marginally higher AUPRC scores with fewer outliers, they suffer from inflated FDR, making their detection results less reliable in practice (Supplementary Fig. S2B, “Outliers”). Among the methods that control FDR, QRscore-Mean performs the best in terms of average AUPRC and detection power.

QRscore-Mean outperforms other methods in differential gene expression analysis when there are excess zeros (Setting 7-9 in Supplementary Table S2). This superior performance is due to its robustness in handling data distribution irregularities, particularly those caused by excess zeros. While random zeros do not impact the direction of mean shifts, they reduce signal strength without inflating false discoveries. This reduction in signal weakens the power of models like edgeR and DESeq2, which do not account for zero inflation.

#### 2.2.4 Three-Sample Tests Show Superiority compared to pair-wise two-sample test in Diverse Simulations for QRscore-Var

We investigated the performance of the three-sample QRscore test compared to the commonly used approach of sequential pairwise testing followed by *p*-value aggregation. In practice, the user may prefer to test the heterogeneity of gene expression across multiple groups or conditions and obtain a single, well-calibrated *p*-value for each gene. Results from Fig. 3 demonstrate that the three-sample QRscore test outperforms the pairwise two-sample tests in terms of AUPRC, particularly when the sample sizes for different groups are not equal. Additional simulations explore scenarios where Groups 1 and 2 were identical with fold changes in Group 3, and cases with varying degrees of dispersion among all three groups. These simulations, which also include the presence of random outliers and zeros, show that the three-sample tests consistently provide superior AUPRC results, demonstrating better adaptability and precision even under model perturbations (Supplementary Fig. S6. Based on the simulation results, we will apply 3-sample tests to GTEx data in Section 2.3.

**Figure 3.**
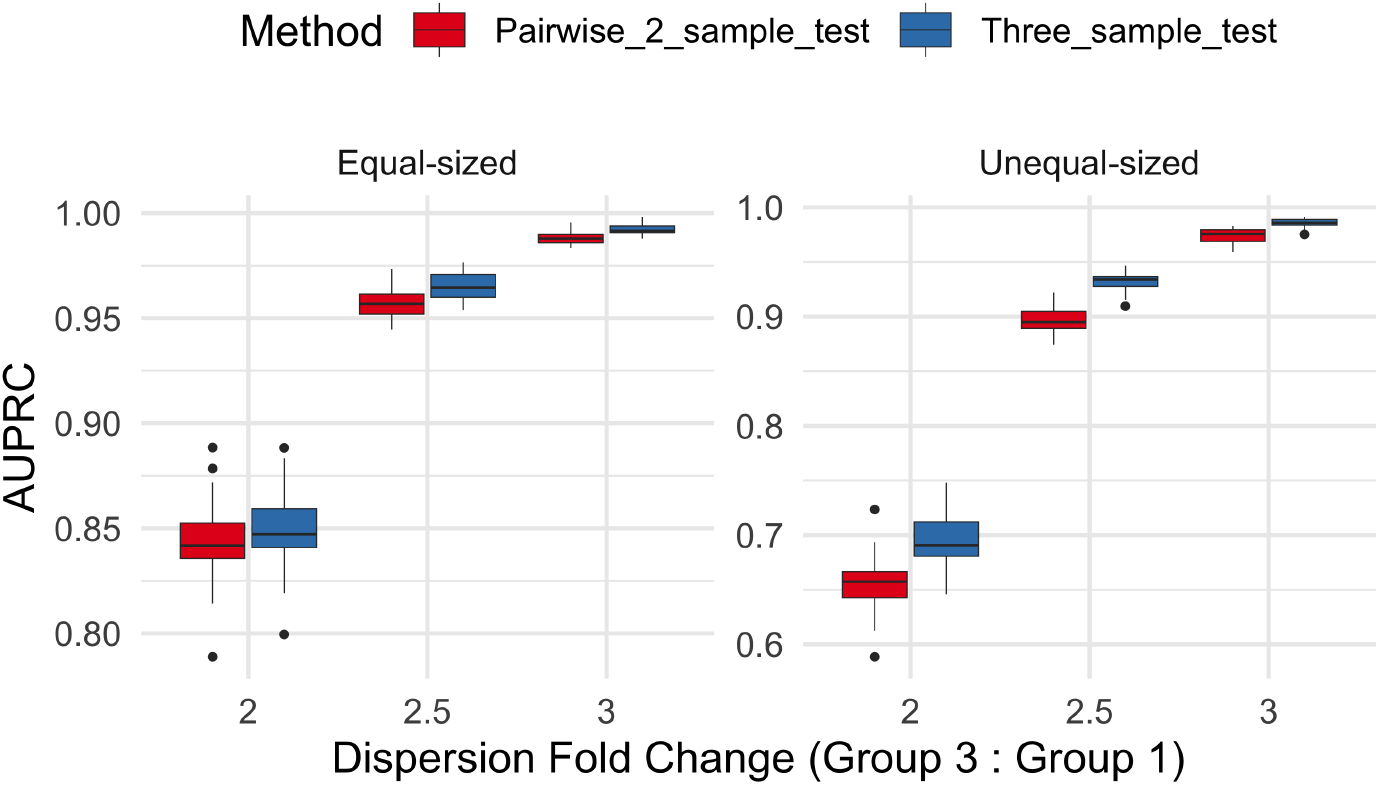
The figure presents two box plots of AUPRC values for the pairwise two-sample test and three-sample test using QRscore-Var. These plots depict conditions with unequal-sized sample sizes (*n*_1_ = 50, *n*_2_ = 100, *n*_3_ = 150) and equal-sized sample sizes (*n*_1_ = *n*_2_ = *n*_3_ = 100), where the dispersion fold change between Group 2 and Group 1 in true DDGs is set to 1:1.5 (all three samples have different dispersions). No zero inflation and outliers are simulated (*α* = *β* = 0).

### 2.3 Application to GTEx

Having demonstrated QRscore’s error control capabilities and enhanced power, we now apply it to the GTEx dataset. The GTEx Project provides RNA sequencing data from bulk tissue samples, gathered from nearly 1,000 deceased donors across various age groups and both sexes. This dataset covers a wide array of tissue types, enabling tissue-specific gene expression analyses.

The flexibility of QRscore allows for the detection of differentially dispersed genes (Section 2.3.1), as well as distinct gene sets that differ in mean expression across tissues (Section 2.3.2). We also show that these gene sets exhibit unique functional effects (Section 2.3.3).

#### 2.3.1 QRscore-Var detects tissue-specific genes with changes in dispersion

We apply QRscore-Var across 33 tissues to identify differentially dispersed genes (DDGs) across three age groups: 20-39 (younger), 40-59 (middle), and 60-79 (older). We identified differing numbers of DDGs across various tissues and detected distinct trends in variance shifts from younger to older age groups (Fig. 4A). In tibial artery and skeletal muscle tissues, most DDGs show an increase in variance from the middle to older age group. In contrast, in whole blood tissue, we observe more DDGs showing an increase in variance from the younger to the middle age group, followed by a decrease from middle to older. Our latter finding agrees with a recent study on gene expression in blood samples from twins, which reported a general decline in expression variability with age [6]. In total, 10,231 unique tissue-specific DDGs were detected. Fourteen tissues reported over 30 DDGs, with whole blood, tibial artery, and transverse colon each reporting more than 2,500 DDGs.

**Figure 4.**
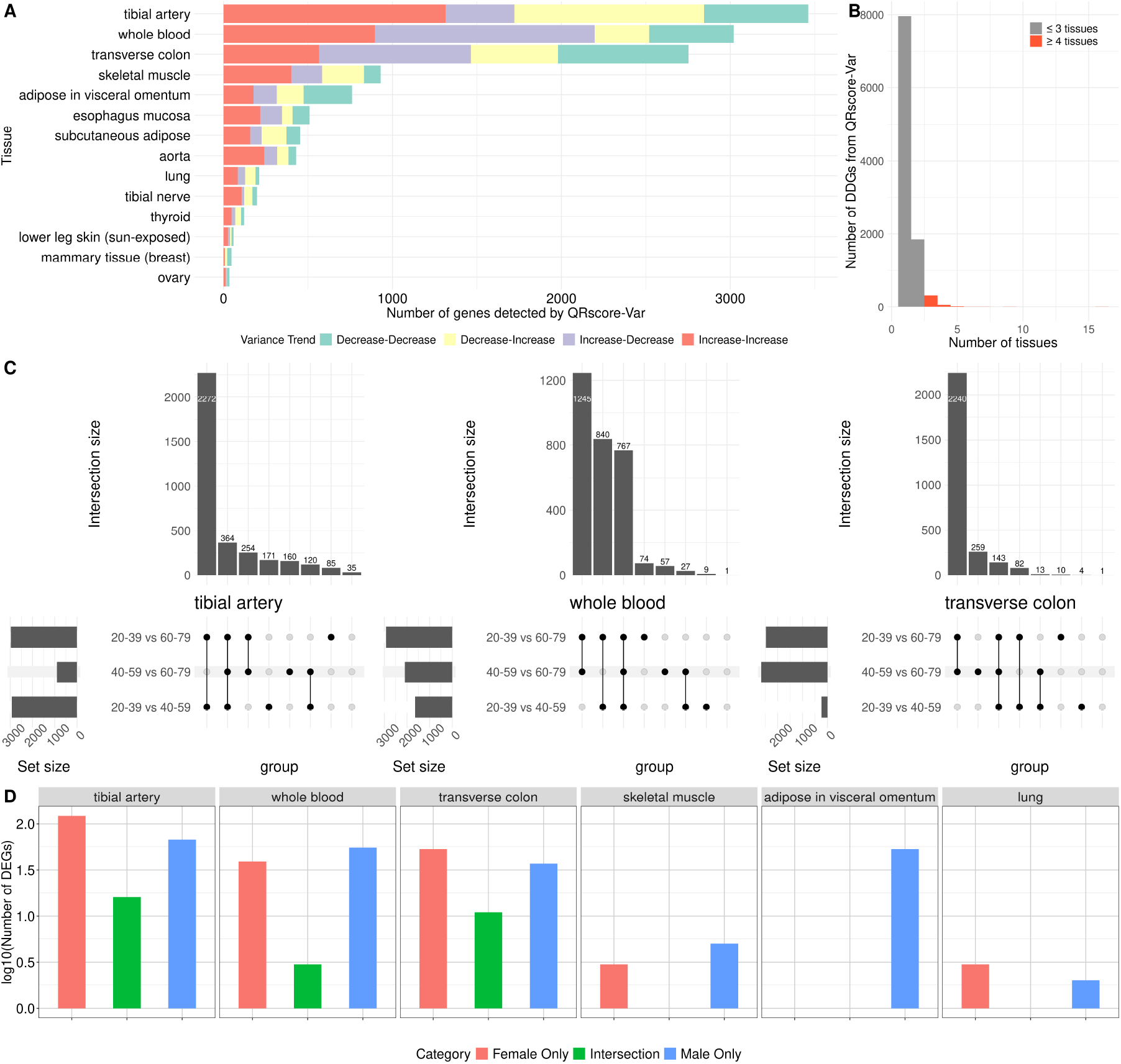
QRscore-Var results on GTEx data.(**A**) A stacked bar plot illustrates the quantity of genes identified by QRscore-Var, categorized by variance trends. The “Decrease-Decrease” label indicates a progressive reduction in gene expression variance from the youngest age group (20-39) through the middle age group (40-59) and into the oldest age group (60-79). Other labels follow a similar naming convention to represent their respective variance trends. (**B**) Histogram of the number of tissues in which each DDG detected only by QRscore-Var is found. The red bars indicate DDGs found from more than 3 tissues. (**C**) Upset barplots illustrates the overlap and distribution of DDGs identified from a three-sample test, further examined through pairwise two-sample tests. This visualization categorizes the DDGs based on their significance in specific pairwise comparisons and the variance shifts between age groups.(**D**) Bar plots display the log counts of DDGs identified in various tissues from female and subsampled male individuals, respectively.

After identifying statistically significant DDGs using the three-sample test, we further characterize the variance shifts between specific age groups by applying pairwise two-sample tests (Fig. 4C). In tibial artery, we found the majority of DDGs are driven by variance shifts between the 20-39 vs. 40-59 and 20-39 vs. 60-79 comparisons, indicating that much of the signal in the three-sample test comes from the younger age group. In transverse colon, the two-sample tests reveal that more DDGs show significant variance shifts between the 40-59 vs. 60-79 and 20-39 vs. 60-79 comparisons, suggesting that the oldest age group exhibits more marked and distinct variation shifts. The detection of more DDGs between the 20-39 and 60-79 age groups is expected, as the broader age range likely leads to more significant differences.

The majority of DDGs are tissue-specific, where each gene was detected in only one tissue (Fig. 4B). However, a few genes show differential dispersion across multiple tissues (Supplementary Table S3). For example, EDA2R, a gene previously known to be associated with aging in the lung [21] and skeletal muscle [22], was detected as a DDG in 16 tissues. Additionally, PTCHD4, previously associated with aging in the thyroid gland [23], was found to have differential dispersion in 16 tissues.

Sex can significantly influence the detection of variance shifts, and the GTEx dataset includes more male than female samples. To address this potential confounding factor, we stratified the data by sex and subsampled male samples to match the number of females, aiming for a fair comparison (Fig. 4D). In tissues such as tibial artery, transverse colon, and whole blood, the numbers of DDGs identified in male and female samples were similar and there are sex-specific DDGs discovered in each tissue. Interestingly, within adipose visceral tissue, even after subsampling to equalize sample sizes, DDGs were identified in male samples but not in female samples. If we do not subsample the data, more DDGs are discovered in males than in females across all tissues that are present in both sexes, due to the larger number of male individuals. This underscores the importance of larger sample sizes for enhancing detection power in DDG analysis, as shown in Supplementary Fig. S7.

#### 2.3.2 QRscore-Var identifies different sets of genes from QRscore-Mean

In parallel with QRscore-Mean, designed to identify genes with mean expression changes using tailored weights, we used QRscore-Var for its capacity to detect variance shifts. Although QRscore-Var detects fewer genes overall compared to QRscore-Mean, it identifies 2,609 unique DDGs across at least one tissue that QRscore-Mean does not capture (that is, DDGs that are not also DEGs; see Fig. 5A), indicating its ability to uncover different aspects of gene expression changes.

**Figure 5.**
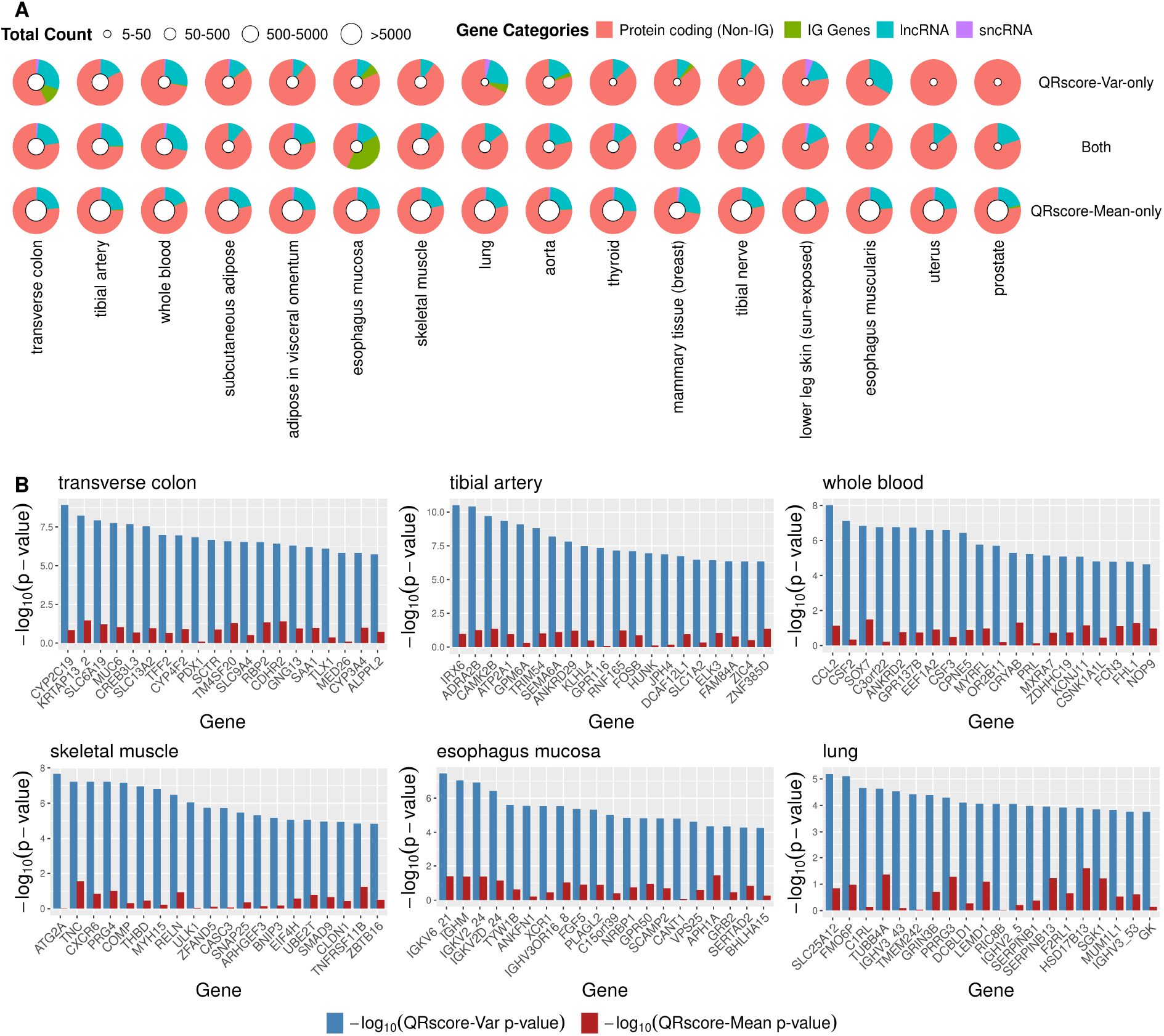
Analysis of Age-related Differential Gene Expression Across Human Tissues Using QRscore-Var and QRscore-Mean. (**A**) Pie charts visualizes the proportion of gene biotypes across various tissues, analyzed through the QRscore-Var and QRscore-Mean methods. The slices represent protein-coding genes, long noncoding RNAs (lncRNAs), and small noncoding RNAs (sncR-NAs), with immunoglobulin (IG) genes separated into their own category due to their substantial proportion in some tissues. Pseudogenes are excluded as they are generally considered non-functional. A white circle at the center indicates the total count of genes. The charts highlight DEGs identified by QRscore-Mean, DDGs identified by QRscore-Var, and genes identified by both methods. Tissues are ranked based on the count of DDGs uniquely identified by QRscore-Var. Bar plots for selected tissues—whole blood, tibial artery, transverse colon, skeletal muscle, lung, esophagus mucosa, and thyroid—highlighting the top 20 significant genes as determined by QRscore-Var. These plots juxtapose the significance levels (− log_10_(*p*-value)) of identified genes between the QRscore-Var and QRscore-Mean analyses, with a focus on protein-coding genes to underline the biological implications.

We found a notable number of DDGs that were identified only by QRscore-Var, but not QRscore-Mean, in whole blood, transverse colon, tibial artery, lung, and thyroid tissues, aligning with studies emphasizing variance shifts in the transverse colon and thyroid [24]. Most DDGs and DEGs are protein-coding genes, with a substantial portion of DDGs comprising immunoglobulin (IG) genes in certain tissues (Fig. 5A). In the esophagus mucosa, IG genes constitute a large proportion of the genes discovered by both QRscore-Var and QRscore-Mean. IG genes are also present in DDGs identified by QRscore-Var in transverse colon and lung, suggesting that they may play a role in immune response alterations associated with aging, as reflected by changes in variability. The discovery of genes from DDG analysis indicates that while mean expression changes reflect overt alterations, variance shifts reveal subtle yet crucial biological variations. This highlights the complex, multifaceted nature of aging at the molecular level, where QRscore-Var captures the nuanced, dynamic alterations that mean-focused analyses might miss.

Fig. 5B illustrates the differential variance expression of the top 20 protein-coding genes across six tissues with aging, as indicated by − log_10_ *p*-values, without shifts in mean expression. Variance shifts in genes like CCL2 in whole blood and SAA1 in transverse colontissues underscore their roles in shaping the inflammatory landscape during aging. CCL2 is pivotal in chronic immune responses during aging [25], while SAA1, contributes to an inflammatory microenvironment that varies across individuals, leading to diverse impacts on epithelial cell function via MMP9 upregulation in aging tissues [25]. In the tibial artery, ADRA2B regulates vascular tone, with its variance shifts suggesting its importance in maintaining vascular health with age. BNIP3 in skeletal muscle indicates diverse mitochondrial balance and autophagy responses, signaling varied susceptibility to age-related muscle degradation [26]. Meanwhile, SERPINB1 in the lung safeguards against protease-induced tissue damage, with its variance shifts revealing intricate age-related changes that impact lung maintenance and contribute to chronic lung conditions [27]. These gene expression shifts reveal the intricate, tissue-specific regulatory changes that contribute to aging and its associated dysfunctions.

To differentiate between genes detected exclusively by either QRscore-Var or QRscore-Mean, we compared their coefficients of variation (CV) ratios and mean ratios across age groups. The CV, a measure of relative variability to the mean, helps to assess data dispersion. Our analysis revealed that genes detected by QRscore-Mean exhibited higher mean changes, whereas genes identified by QRscore-Var showed greater changes in CV (Supplementary Fig. S8).

We compiled lists of genes related to human aging from three databases: GenAge [28], LongevityMap [29], and CellAge [30] to annotate age-related genes. We ranked the genes based on *p*-values for each tissue and age group, using QRscore-Var or QRscore-Mean tests, and applied one-sided Wilcoxon rank-sum tests to evaluate the methods’ effectiveness at prioritizing aging-related genes. Interestingly, in Table 3, QRscore-Var showed statistical significance (*p <* 0.05) in ranking aging-related genes in twelve out of fourteen tissues. However, the Wilcoxon test *p*-values were less significant than QRscore-Mean in most tissues and not significant in lung and lower leg skin (sun-exposed). This suggests that, as the existing databases are mostly annotated by mean expression shifts, they may miss age-related DDGs that QRscore-Var can identify. In doing so, QRscore-Var provides a complementary perspective on the genetic basis of aging.

**Table 3:** *p*-values from one-sided Wilcoxon rank-sum tests for ranking aging-related genes above other genes by QRscore-Var or QRscore-Mean. The tissues in the table have at least one age-related gene discovered only by QRscore-Var.

| Tissue | QRscore-Var | QRscore-Mean |
| --- | --- | --- |
| tibial nerve | $1.3 \times 10^{-7}$ | $2.0 \times 10^{-9}$ |
| skeletal muscle | $2.0 \times 10^{-7}$ | $1.8 \times 10^{-2}$ |
| esophagus muscularis | $6.5 \times 10^{-6}$ | $3.2 \times 10^{-5}$ |
| subcutaneous adipose | $1.2 \times 10^{-5}$ | $5.8 \times 10^{-9}$ |
| thyroid | $5.6 \times 10^{-4}$ | $3.5 \times 10^{-5}$ |
| tibial artery | $2.5 \times 10^{-3}$ | $7.3 \times 10^{-5}$ |
| mammary gland (breast) | $4.5 \times 10^{-3}$ | $2.6 \times 10^{-4}$ |
| adipose in visceral omentum | $1.2 \times 10^{-2}$ | $2.2 \times 10^{-6}$ |
| whole blood | $2.2 \times 10^{-2}$ | $2.1 \times 10^{-3}$ |
| transverse colon | $2.9 \times 10^{-2}$ | $1.5 \times 10^{-10}$ |
| aorta | $4.0 \times 10^{-2}$ | $2.6 \times 10^{-5}$ |
| esophagus mucosa | $4.2 \times 10^{-2}$ | $2.4 \times 10^{-5}$ |
| lower leg skin (sun-exposed) | $1.5 \times 10^{-1}$ | $1.2 \times 10^{-5}$ |
| lung | $1.6 \times 10^{-1}$ | $6.6 \times 10^{-14}$ |

#### 2.3.3 QRscore-Var and QRscore-Mean identify genes that are functionally distinct

To further investigate the types of genes identified by QRscore-Var and QRscore-Mean, we perform Gene Ontology (GO) enrichment analysis. We looked for enriched terms in biological process among DDGs and DEGs detected by QRscore.

QRscore-Var identifies a substantial number of DDGs in whole blood tissue. To gain deeper insights into these genes and their gene ontology, we examined the top 10 enriched terms in biological process for top 500 significant genes ranked by QRscore-Var and QRscore-Mean (Fig. 6A&B). While QRscore-Mean highlights fundamental processes such as mRNA processing and RNA splicing, QRscore-Var uncovers additional layers of complexity in gene expression patterns related to angiogenesis, positive regulation of the ERK1/2 cascade, and various types of chemotaxis. Enriched ontology terms of DDGs in whole blood particularly highlighted the dynamic regulation of cellular responses, emphasizing processes such as angiogenesis and immune cell migration, which are crucial for responding and adapting to physiological changes. Variance analysis reveals a broader spectrum of gene expression responses, showing that the robustness of cellular processes, like cascade regulation and cell migration, goes beyond changes in average expression.

**Figure 6.**
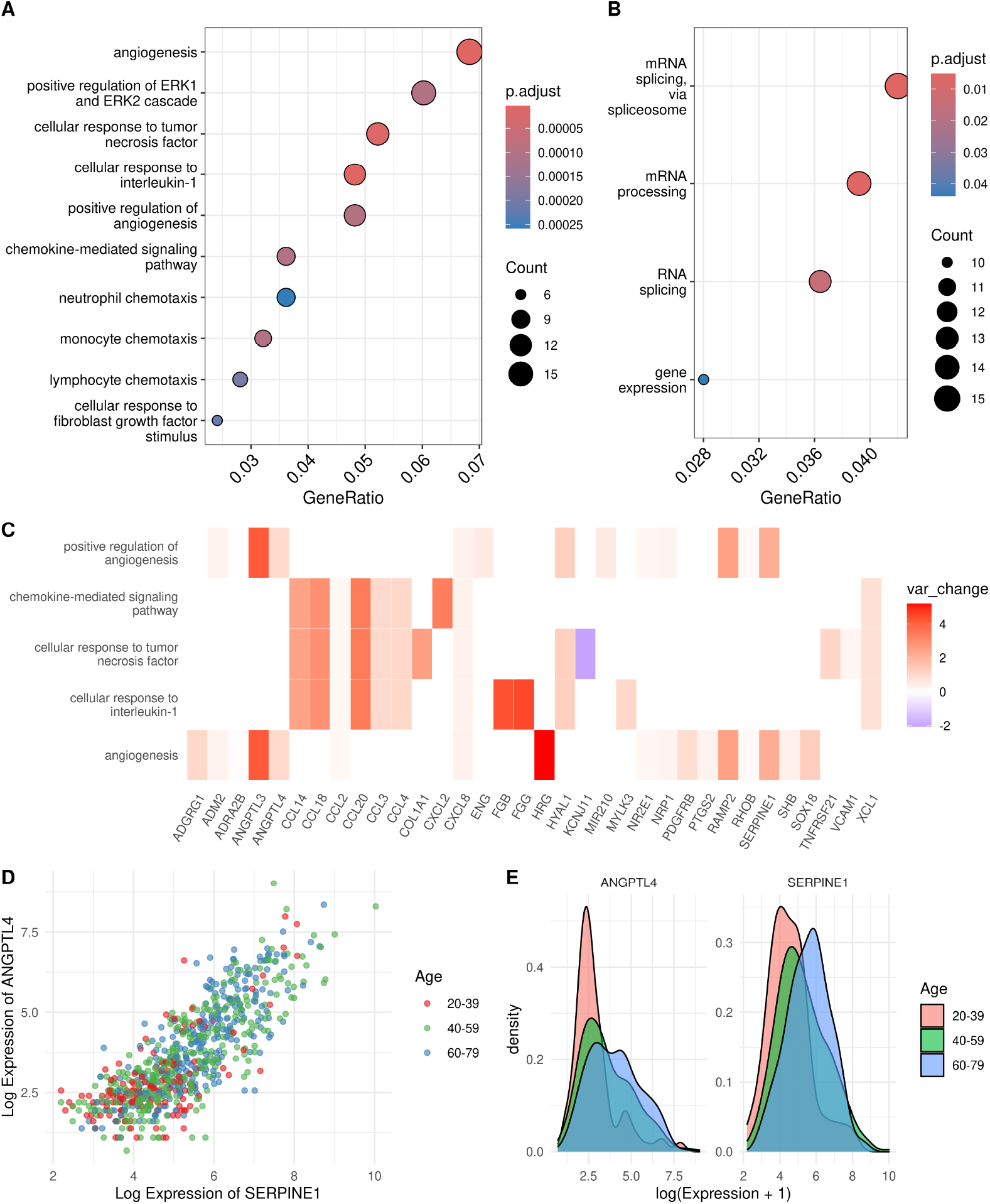
Comparative analysis of GO enrichment in whole blood tissue based on significant genes identified by QRscore-Var and QRscore-Mean. (**A**) A dot plot illustrates the top 10 enriched GO terms within the biological process ontology for top 500 genes significantly detected by QRscore-Var. (**B**) Similar enrichment analysis results for genes identified by QRscore-Mean. **(C)** A heat plot, generated using ClusterProfiler, shows the fold change in expression variance between the 20-39 and 40-59 age groups for significant genes associated with the top 5 enriched GO terms identified by the QRscore-Var analysis. **(D)** A dot plot and **(E)** a density plot further investigate the expression patterns of two highly coexpressed genes identified by QRscore-Var as being involved in angiogenesis.

In Fig. 6C, we observe large fold changes in variance for genes such as HRG, ANGPTL3, and SERPINE1, which are closely linked to angiogenesis, as well as for FGB, and FGG, which play key roles in the cellular response to interleukin-1 and regulation of blood coagulation. Additionally, the analysis reveals notable variance shift among age groups in the C-C motif chemokine ligand (CCL) gene family, such as CCL14, CCL18, and CCL20, which are enriched in chemokine-mediated signaling pathways and cellular response to tumor necrosis factor. This further highlights the intricate gene expression dynamics involved in immune responses and inflammation.

Furthermore, Table 4 offers a detailed comparison between QRscore-Var and QRscore-Mean analyses, highlighting the top 20 genes that contribute to the ontology terms in Fig. 6A for DDGs identified by QRscore-Var in whole blood tissue. Notably, CCL2 is uniquely identified as significant by QRscore-Var, with 18 additional genes ranked in the top 200 by QRscore-Var but outside the top 1000 for QRscore-Mean. CCL2 exhibits increasing dispersion from young to middle age, followed by a decrease from middle to old age, revealing complex regulatory dynamics in disease pathogenesis (Supplementary Table S5). This highlights the unique insights provided by analyzing gene expression variability. Moreover, we identified pairs of DDGs with highly correlated gene expression that are key contributors to each enriched GO term. One example is shown in Fig. 6D&E, where SERPINE1 and ANGPTL4, both identified as DDGs, are coexpressed with a Spearman’s correlation of 0.808 and are implicated in the regulation of angiogenesis and blood coagulation. Details of all coexpressed DDG pairs for each tissue are provided in Supplementary Table S4. This coexpression analysis, along with the variance and GO enrichment findings, illustrates the nuanced interplay of genes in aging, demonstrating the value of integrating QRscore-Var and QRscore-Mean analyses to uncover the broad spectrum of biological processes affected by aging in human tissues. Additionally, across diverse tissues including tibial artery, transverse colon, skeletal muscle, and thyroid, QRscore-Var analysis has illuminated key biological processes and genes that exhibit significant variance in expression with aging. In the tibial artery, variance shifts in DDGs highlight tissue-specific processes such as sprouting angiogenesis and the positive regulation of cell migration, both of which are crucial for maintaining vascular integrity and promoting repair. This emphasis on angiogenesis, also observed in whole blood, underscores the artery’s significant role in these processes (Supplementary Fig. S9). Beyond these processes, SEMA6A, identified as a DDG but not a DEG, is associated with the semaphorin-plexin signaling pathway and the negative regulation of axon extension involved in axon guidance, pointing to additional molecular mechanisms at play in the tibial artery (Supplementary Table S5). In the transverse colon, DDGs involved in maintaining the gastrointestinal epithelium and lipid metabolism are critical for preserving structural integrity and supporting digestion (Supplementary Fig. S10). In skeletal muscle (Supplementary Fig. S11), coexpressed DDGs like COMP and FMOD underscore their importance in collagen fibril organization and maintaining the structural integrity of muscle tissue. Moreover, several Protocadherin Gamma Subfamily genes are identified as DDGs, such as PCDHGA10 and PCDHGB2, and they are linked to nervous system development and was found to be associated with muscle weakness during aging [31]. In the thyroid gland, DDGs involved in the hydrogen peroxide catabolic process and cellular oxidant detoxification are essential for regulating hormone synthesis and preventing oxidative damage during aging. The thyroid’s role in immune response and its adaptation to metabolic demands are further highlighted by DDGs in the stimulatory C-type lectin receptor signaling pathway, particularly with the coexpression of KLRC4 and KLRK1 (Supplementary Fig. S12). This concise synthesis across varied tissues not only pinpoints specific genes and biological processes influenced by aging but also enhances our understanding of aging’s tissue-specific effects, offering valuable directions for future research and intervention strategies to combat aging’s adverse outcomes.

**Table 4:** Results of QRscore-Var and QRscore-Mean for the top 20 genes ranked by QRscore-Var on three age groups in whole blood tissue, considering only genes that are in the top 10 enriched GO terms in biological process from DDGs detected by QRscore-Var (Fig. 6A).

| Gene | QRscore-Var |  | QRscore-Mean |  |
| --- | --- | --- | --- | --- |
| | Adjusted $p$-value | rank | Adjusted $p$-value | rank |
| CCL20 | $6.2 \times 10^{-10}$ | 8 | $4.8 \times 10^{-3}$ | 11284 |
| EGR3 | $7.6 \times 10^{-10}$ | 9 | $9.8 \times 10^{-4}$ | 9673 |
| HRG | $1.8 \times 10^{-8}$ | 16 | $3.9 \times 10^{-4}$ | 8883 |
| NR4A1 | $1.9 \times 10^{-8}$ | 18 | $4.5 \times 10^{-5}$ | 7193 |
| CXCL8 | $1.6 \times 10^{-7}$ | 36 | $1.3 \times 10^{-2}$ | 12554 |
| RHOB | $1.6 \times 10^{-7}$ | 40 | $6.0 \times 10^{-11}$ | 1203 |
| VCAM1 | $3.1 \times 10^{-7}$ | 50 | $3.7 \times 10^{-9}$ | 2383 |
| MIR210 | $3.5 \times 10^{-7}$ | 52 | $2.9 \times 10^{-7}$ | 4261 |
| CCL14 | $6.8 \times 10^{-7}$ | 67 | $9.5 \times 10^{-6}$ | 6207 |
| CCL18 | $7.2 \times 10^{-7}$ | 69 | $7.4 \times 10^{-4}$ | 9410 |
| FGG | $7.7 \times 10^{-7}$ | 75 | $6.2 \times 10^{-6}$ | 5939 |
| FGA | $9.9 \times 10^{-7}$ | 78 | $9.9 \times 10^{-6}$ | 6234 |
| NR2E1 | $9.9 \times 10^{-7}$ | 79 | $6.9 \times 10^{-4}$ | 9340 |
| SERPINE1 | $2.1 \times 10^{-6}$ | 98 | $8.4 \times 10^{-6}$ | 6125 |
| CCL2 | $2.3 \times 10^{-6}$ | 99 | $1.1 \times 10^{-1}$ | 16110 |
| CXCL2 | $3.3 \times 10^{-6}$ | 110 | $1.0 \times 10^{-4}$ | 7801 |
| APOE | $5.2 \times 10^{-6}$ | 125 | $1.7 \times 10^{-6}$ | 5162 |
| CCL3 | $6.9 \times 10^{-6}$ | 142 | $2.7 \times 10^{-12}$ | 587 |
| FGB | $6.9 \times 10^{-6}$ | 145 | $4.1 \times 10^{-5}$ | 7126 |

## 3 Discussion

We introduced QRscore, a suite of rank-based non-parametric tests for identifying DDGs and DEGs from bulk RNA-seq data. We showed, through extensive simulations, that QRscore is robust at detecting both mean and dispersion shifts, particularly under challenging and realistic conditions where datasets have inflated zeros and outliers. Notably, QRscore controls false discovery well and achieves high power even in such settings. Applying QRscore to GTEx data, we find different DDGs and DEGs in each tissue that are also functionally distinct, implicating novel candidates for further biological investigation.

Model misspecification is a critical issue in RNA-seq data analysis that potentially explains the inflation of false discoveries observed in practice. For instance, a study [32] has shown that methods relying on specific distributional assumptions, like NB or ZINB, identify different sets of DEGs when applied to RNA-seq data from different mouse strains and human tissues, with their FDR significantly increasing in the presence of outliers. QRscore, as a rank-based non-parametric method, is less sensitive to underlying data distribution and effectively handles datasets with technical zeros and outliers. Thus, being less prone to model misspecification, QRscore delivers more reliable and replicable results that correspond to likely true biological signals. This would potentially improve the true discovery rate of experiments downstream of gene prioritization, such as functional assaying.

As pointed out in the Introduction, gene expression variability plays an important role in aging, adaptation to changing environments, and disease susceptibility. Focusing on aging in our present work, senescence-related changes in gene expression variability are often driven by shifting regulatory efficiency and the accumulation of epigenetic modifications [33], which may go unnoticed when focusing solely on mean expression levels. QRscore addresses this gap by detecting variance shifts, uncovering previously unreported age-related DDGs. For instance, in whole blood, we find links between DDGs and angiogenesis, a process not prominently highlighted by DEGs. Similarly, SEMA6A was identified as a DDG in tibial artery tissue. Despite not being classified as a DEG in this tissue and previously found not to be associated with age in tumor tissues [34], this finding suggests SEMA6A’s potential connection to aging may have been overlooked owing to a focus on mean shifts. These observations underscore the importance of analyzing variance shifts and other distributional changes to uncover biologically relevant insights, not only in aging but also under broader conditions.

Our present work is not without limitations. While QRscore effectively controls Type I error, it tends to be conservative and requires larger sample sizes to ensure reliable results — a constraint inherent to rank-based tests. Fortunately, in recent years many scholars have emphasized the importance of using larger datasets when studying differential variability or distribution. Building on our current findings with sufficiently large samples like GTEx, future work will focus on applying QRscore to even larger and more diverse datasets. Additionally, QRscore can be adapted to accommodate different parametric distributions and target other types of distributional changes, potentially guided by biologically informed hypotheses. This could potentially lead to new insights into gene expression dynamics.

The future of differential expression analysis lies in the ability to detect and explain all patterns of change in differential expression, accounting for confounders, technical variation, and noise in the data. Flexible tests like QRscore provide a path to realizing this goal by accommodating various data characteristics and distributional shifts. Our work therefore represents a small but meaningful contribution towards this objective. As tools for DE analysis become more flexible toward types of distributional shifts, biologists can expect to detect and explain more ways in which genes are differentially expressed between and across conditions, thereby providing a more complete understanding of the vast landscape of gene expression across multitudinous contexts.

## 4 Methods

### 4.1 Non-parametric tests for detecting differential mean or differential dispersion in RNA-seq data

We present a rank-based test that capitalizes on the benefits of parametric modeling while remaining non-parametric in nature. This approach inherits the strengths of non-parametric tests, such as false positive control, while also providing increased detection power if the data roughly follow a parametric model.

#### 4.1.1 Weight construction

To obtain the weight functions *w*_*g*_ appearing in Eq. (1), we adopt the approach of [35], which combines likelihood gradients (i.e., scores) with rank statistics to construct *w*_*g*_. Erdmann-Pham showed that for many asymptotically efficient non-parametric tests (e.g., Mann-Whitney, van der Waerden, and Klotz tests) — that is, tests that surprisingly achieve similar power to their parametric counterparts in large samples when the parametric assumptions are met — their weight functions satisfy a first-order Taylor approximation to the difference between the log-likelihood under the null hypothesis *P*_*g*_ = *Q*_*g*_ and the log-likelihood under a “slightly perturbed” alternative hypothesis. Here, *P*_*g*_ and *Q*_*g*_ are parametric distributions for which the non-parametric test is compared against its parametric counterpart, and “slightly perturbed” refers to a very small perturbation of the associated parameter. For example, *P*_*g*_ and *Q*_*g*_ could be normal distributions parameterized by different means *θ*_*P*_ and *θ*_*Q*_, with the van der Waerden test being asymptotically as powerful as the *t*-test when samples themselves are drawn from normal distributions with different means (and the Mann-Whitney test also being comparable to the *t*-test), and the perturbation is 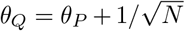 where *N* = *n* + *k* is the pooled sample size.

The asymptotic efficiency for above non-parametric tests, however, is limited to normal distributions. Our method extends this approach by designing weights informed by any parametric model while maintaining a non-parametric framework. In this study, we construct *w*_*g*_ tailored to the (zero inflated) negative binomial family, which better models gene expression data. As we will show, these weights are derived by computing the score of a transformed rank statistic, which handles over-dispersion and zero inflation in RNA-seq data (Methods 4.1.3)

Concretely, for each gene, we model the count *Y*_*jg*_ using a negative binomial distribution *NB* (*µ*_*g*_, *r*_*g*_), where *µ*_*g*_ and *r*_*g*_ are the mean and dispersion parameters, respectively. We then propose *X*_*ig*_ to follow *NB* (*θ*_1_*µ*_*g*_, *θ*_2_*r*_*g*_), with *θ*_1_ and *θ*_2_ representing fold changes in mean and dispersion. Note that this modeling step is not for formulating the null hypothesis but only for constructing the weights.

To target dispersion shifts, we set *θ*_1_ = 1 and define our score function as 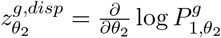, where 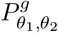 represents the probability distribution of *X*_*ig*_ under the parameters *θ*_1_*µ*_*g*_ and *θ*_2_*r*_*g*_. Given the cumulative distribution function (CDF) denoted as 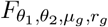,the corresponding weight function is:

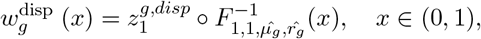

where 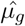 and 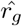 are maximum likelihood estimates of the negative binomial distribution parameters, computed from empirical counts data. The symbol ○ represents function composition, meaning the inverse CDF 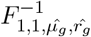 is applied first and its output is passed as input to 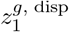.

This weight construction employs a modified Score test, where the original samples are replaced by functions that take in rank-based statistics. Under the null hypothesis that gene expression distributions are identical under two conditions, the ranks 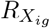 are uniformly distributed in {1, 2, …, *n* + *k*}. In this context, the inverse CDF becomes pivotal, mapping this uniform distribution back into the scale of negative binomial distribution. This step adapts rank-based data for use in the Score test, while maintaining its non-parametric nature.

Crucially, the use of rank-based statistics as inputs guarantees that the test is distribution-free. This ensures robustness against model misspecification and provides stringent control over false discoveries. Equally important is the method’s efficiency: when parameters and the score function are tailored within a negative binomial model, the test’s ability to detect changes in dispersion is remarkably enhanced. Most notably, with an accurately specified model, this test is asymptotically as efficient as that of the likelihood-ratio test. This elevates it above other non-parametric tests in scenarios where the model precisely captures the underlying distribution.

Similarly, we can set *θ*_2_ = 1 and derive the corresponding score function and test statistic 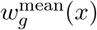 that targets on shifts in mean parameters. A detailed derivation of the functions used in constructing the weights, along with numerical considerations and adjustments, is provided in Appendix A.

#### 4.1.2 Inference

The asymptotic normality and efficiency of the general test statistic are proved in [35]. In the context of this work, suppose *k, n* → ∞, with *k/*(*n* + *k*) → *α >* 0. The statistic 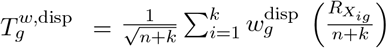 is asymptotically normal under null hypothesis *H*_0*g*_ with

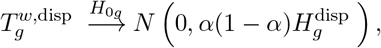

where 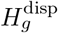 is the Fisher information specific to the dispersion parameter in the NB distribution. For each gene, the *p*-values are obtained using the asymptotic distribution of the above statistics and adjusted for multiple testing using the Benjamini-Hochberg (BH) procedure. We refer to our method as QRscore-Var when assessing dispersion changes and QRscore-Mean when assessing mean changes.

Furthermore, our QRscore framework includes extensions to both zero-inflation models (Methods 4.1.3) and *K*-sample tests (Methods 4.1.4). These extensions allow QRscore to be applied to a wider range of RNA-seq data analysis scenarios, accommodating the presence of technical zeros and multiple sample groups.

#### 4.1.3 Extension to zero-inflation model

Large-scale RNA-seq data often contain a substantial number of genes with technical excess zeros [13], which can pose a challenge for accurate analysis of gene expression means and variability. To properly account for such zeros to ensure reliable and interpretable results, it is customary to model the counts for each gene in bulk RNA-seq data by a zero-inflated negative binomial distribution ZINB(*µ*_*g*_, *r*_*g*_, *π*_*g*_):

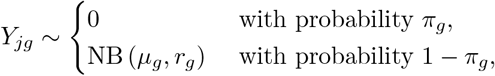

where 0 ≤ *π*_*g*_ *<* 1 is the probability of technical excess zeros. This model facilitates the use of probabilistic methods to distinguish between zero counts resulting from technical variations at the excess zero state and those arising from biological variations at the negative binomial state.

In a manner akin to the weight construction for the NB model, we model *X*_*ig*_ by ZINB (*θ*_1_*µ*_*g*_, *θ*_2_*r*_*g*_, *π*_*g*_).

The score function, CDF, and Hessian values for this model are derived from the ZINB probability mass function. We estimate the parameters *µ*_*g*_, *r*_*g*_, and *π*_*g*_ using numerical maximum likelihood estimates obtained from the pscl package [36].

Since technical zeros are not present in all genes, we fit the negative binomial model and designed weights described in Methods 4.1.1 for genes with expression levels all greater than zero in each sample. For genes containing zeros, we applied the zero-inflated model in further analyses. It’s important to note that if all zeros in a gene’s expression profile originate from the negative binomial model (*π*_*g*_ = 0), the numerical maximum likelihood estimate of *π*_*g*_ would be very close to zero. Consequently, this scenario results in a test that behaves similarly to the negative binomial weights. Alternatively, we can evaluate the presence of excess zeros by testing the hypothesis 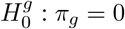 against 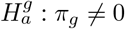 in the zero-inflated model. Under the null hypothesis when *π*_g_ = 0, all of the zero counts are assumed to arise from biological variations at the negative binomial state.

Similar to MDSeq [13], we fit a negative binomial model and a zero-inflated negative binomial model for each gene and then apply a likelihood ratio test between two models. When 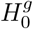 is rejected, the zero-inflated model is applied instead of the negative binomial model for further analyses. This model-selection step ensures that subsequent tests for differential expression among genes, which rely on the appropriate model, yield valid *p*-values (as demonstrated in [35]).

#### 4.4.4 *K*-sample test

We next generalize our procedure to a *K*-sample scenario, where we have *K* distinct samples. Suppose there are K samples for each gene *g*, and our null hypothesis is

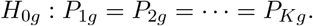

The alternative hypothesis *H*_1*g*_ : at least one distribution is different from the others.

Suppose the sample sizes are *n*_1_, *n*_2_, …, *n*_*k*_. Let 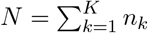 and 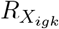 be the rank of ith value in sample k after pooling and sorting all the samples for gene g. The test statistic

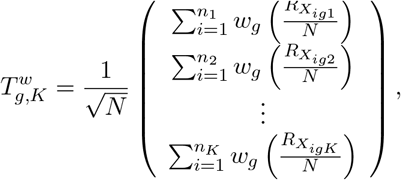

where *w* is constructed the same way as our two-sample test.

The general idea of the *K*-sample test is to efficiently combine information from multiple group comparisons into a unified analysis. For each gene, we isolate one group at a time and get a two-sample test statistic between that group and the pooled remaining groups. This results in a vector of test statistics, one for each group. Importantly, instead of conducting inference separately and obtaining K *p*-values for each gene, we project this vector into a subspace to break the correlation between the group-wise statistics. After this projection, we derive the covariance matrix of the transformed statistics and apply a chi-square test, yielding a single, well-calibrated *p*-value for each gene. The detailed test statistic construction and inference is in Appendix B.

### 4.2 Simulation study

We conduct simulations to evaluate the performance of methods for detecting differential expression driven by changes in either dispersion or mean across groups. Simulations are based on parameters estimated from bulk RNA-seq count matrices from whole blood tissue of individuals aged 40-49 and 60-69 from the GTEx project.

We first estimate the necessary parameters for each gene. We filter out genes with more than 20% zeros, a step implemented only during the setup of the simulation pipeline to prevent the occurrence of inflated zeros unless they are deliberately introduced in the simulation. We then estimate size factors (*Ŝ*_*i*_) for each individual *i* using the median of scaled counts across genes as applied in DESeq2. After normalization of the raw counts using these size factors, we obtained the mean 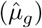 and dispersion 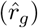 parameters for each gene.

Since there are *n*_1_ = 113 individuals in the 40-49 age group and *n*_2_ = 249 individuals in the 60-69 age group from whole blood tissue, we simulate *N* × *G* count matrices, where *N* = *n*_1_ + *n*_2_ = 382 and *G* = 3000 in both the dispersion shift and mean shift simulations. For the dispersion shifts simulations, we apply specific fold changes to 10% of the genes, randomly selected as true DDGs. In the basic setting, all genes in Group 1 and non-DDGs in Group 2 are simulated using negative binomial models based on estimated parameters (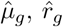and *Ŝ*_*i*_). For the selected DDGs in Group 2, a single fold change value for the dispersion parameter is applied uniformly within each dataset. This value ranges from 1.1 to 2.9 across different simulated datasets, generating varying levels of differential dispersion. We simulate increased and decreased dispersion separately for Group 2 relative to Group 1, allowing us to assess the impact of these changes on detection methods. To further test robustness, we vary sample sizes *n*_1_ and *n*_2_ and introduce scenarios with different proportions of genes that contain outliers or inflated zeros (Table 5). Similarly, for mean shift simulations, we apply fold changes to 10% of the genes, randomly selected as true DEGs, with values ranging from 1.1 to 2.0, incremented by 0.1, and the direction of the shift is randomized. We repeat each simulation across 20 random seeds to account for variability and evaluate the stability of the methods under different conditions. The details of the simulation is described in Appendix C. To evaluate the type I error in decreased sample sizes, we simulated data from purely negative binomial model without any fold change spike-ins and any distribution perturbations. The simulations maintain balanced sample sizes between two groups, varying from 10 to 100.

**Table 5:** Simulation settings. In the basic simulation, setting 1 simulates data from negative binomial distributions (*n*_1_ = 113, *n*_2_ = 249). Settings 2 - 3 follow the same approach as Setting 1 but with smaller sample sizes of *n*_1_ = 50, *n*_2_ = 100 and *n*_1_ = *n*_2_ = 50. Settings 4 to 10 have the same sample sizes as Setting 1. Settings 4 - 6 introduce random outliers in 5%, 10%, and 20% of the genes. Settings 7 - 9 simulate random zeros in 5%, 10%, and 20% of the genes. Setting 10 simulate random outliers in 10% genes and random zeros in 10% genes. Settings 11 and 12 replicate Setting 10 with smaller sample sizes of *n*_1_ = 50, *n*_2_ = 100 and *n*_1_ = *n*_2_ = 50.

| Setting | $n_1$ | $n_2$ | Proportion of genes<br>with outliers ($\alpha$) | Proportion of genes<br>with inflated zeros ($\beta$) |
| --- | --- | --- | --- | --- |
| 1 | 113 | 249 | 0 | 0 |
| 2 | 50 | 100 | 0 | 0 |
| 3 | 50 | 50 | 0 | 0 |
| 4 | 113 | 249 | 5% | 0 |
| 5 | 113 | 249 | 10% | 0 |
| 6 | 113 | 249 | 20% | 0 |
| 7 | 113 | 249 | 0 | 5% |
| 8 | 113 | 249 | 0 | 10% |
| 9 | 113 | 249 | 0 | 20% |
| 10 | 113 | 249 | 10% | 10% |
| 11 | 50 | 100 | 10% | 10% |
| 12 | 50 | 50 | 10% | 10% |

Finally, to compare the performance of three-sample QRscore-Var tests with sequential pairwise two-sample tests, we simulate gene expression data for both equal sized (*n*_1_ = *n*_2_ = *n*_3_ = 100) and unequal sized samples (*n*_1_ = 50, *n*_2_ = 100, and *n*_3_ = 150). We randomly select 10% of the genes to be true DDGs. We assess heterogeneity across multiple groups, evaluating two setups: one with identical distributions for Groups 1 and 2 and fold changes in Group 3, where the dispersion fold change between Group 3 and Group 1 is set to 1.5, 2, and 2.5; and another with differences in all three groups, specifically, dispersion fold change from Group 2 to Group 1 is 1.5, and from Group 3 to Group 2 is 2, 2.5, and 3. We simulate the data using pure negative binomial distributions or randomly selected 10% genes with outliers and 10% genes with random zeros, incorporating fold changes for selected true DDGs.

### 4.3 Real data analysis

In real data applications, we use publicly available bulk RNA-seq data across multiple human tissues provided by the Genotype-Tissue Expression (GTEx) Consortium [19].

We divide bulk RNA-seq count data by three age groups, namely 20-39, 40-59 and 60-79. We selected 33 tissues, each containing more than 25 donors, for differential analysis. We also filtered out lowly expressed genes, defined as those with an average count of less than 2 and more than 80% zeros, resulting in approximately 50% of genes in each group being retained for further analysis (Supplementary Fig. S13). Before differential analysis, we apply a normalization procedure provided by DESeq2, which divides counts by sample-specific size factors determined by median ratio of gene counts relative to geometric mean per gene.

We apply QRscore on GTEx data after prefiltering and normalizing the data. Using QRscore-Var and QRscore-Mean, we conduct three-sample tests on three age groups for each tissue, adjusting the *p*-values with BH adjustments to identify sets of DDGs and DEGs (threshold 0.05). For further insights, we calculate the variance ratio for pairwise two-samples and performed pairwise two-sample tests on selected DDGs to determine which groups drive the signal. Additionally, we separate individuals by sex and perform the same tests for each sex group respectively. We also conduct the tests on subsampled male data to match the sample size with females for a fair comparison.

For futher annotation, we compile lists of genes related to human aging from three databases: GenAge [28], LongevityMap [29], and CellAge [30] to annotate age-related genes. We rank the genes based on *p*-values for each tissue and age group, using QRscore-Var or QRscore-Mean tests, and applied one-sided Wilcoxon rank-sum tests for the age-related genes against other genes. We conduct ontology enrichment analysis using the ClusterProfiler package. For coexpressed gene analysis, we extracted pairs of DDGs enriched in each GO term and calculated their Spearman correlation. Gene pairs with an absolute correlation greater than 0.6 were retained.

### 4.4 Other methods for identifying DDGs and DEGs

We utilized various other methods for benchmarking, including MDSeq, GAMLSS, diffVar, Levene’s test, and Bartlett’s test for detecting DDGs, as well as DESeq2, edgeR, limma-voom, and the Mann-Whitney test for detecting DEGs. All these methods took a read count matrix and a condition label vector as input. The parameters were set based on the user guides of these methods’ software packages. We applied Benjamini-Hochberg Procedure for *p*-values to control FDR.

For diffVar, we first applied the normalization procedure in edgeR and then used varFit function from package missMethyl (v11.32.0) to perform differential analysis.

For MDSeq (v1.0.5), we first calculated normalization factors from edgeR package, then applied MDSeq function and set normalization factors as an offset, followed by generating the results using the extract.ZIMD function.

For GAMLSS, we calculated the normalization factor in the same way as we did for MDSeq. Then we applied function gamlss in gamlss package (v5.4-12) with and without regression model targeting on dispersion parameter added and perform likelihood ratio tests.

For Levene’s test, we normalized the data the same way as DESeq2, and then we did log transformation for each gene to increase detection power. We used leveneTest in package car (v3.1-1) to optain *p*-value for each gene.

For edgeR (v4.2.1), DESeq2 (v1.44.0) and limma (v3.60.3), we used their default parameters for normalization and DEG analysis.

## 5 Availability of data and materials

The GTEx data are available from the GTEx Portal: https://storage.googleapis.com/adult-gtex/bulk-gex/v8/rna-seq/GTEx_Analysis_2017-06-05_v8_RNASeQCv1.1.9_gene_reads.gct.gz. The age-related gene annotations are from GenAge: https://genomics.senescence.info/genes/human_genes.zip; LongevityMap: https://genomics.senescence.info/longevity/longevity_genes.zip and CellAge: https://genomics.senescence.info/cells/cellAge.zip

## 6 Code availability

The R package for QRscore is available at https://github.com/songlab-cal/QRscore.

## Acknowledgments

This research is supported in part by an NIH grant R35-GM134922 and grant number CZF2019-002449 from the Chan Zuckerberg Initiative Foundation.

## Appendix A Test Statistic Details

### A.1 Formula Derivation for Test Statistic in QRscore-Var

To derive the quantities used to compute the test statistic targeting the dispersion in zero-inflated negative binomial distribution (ZINB), we first consider a negative-binomial distribution, *NB*(*θ*_1_*µ*_*g*_, *θ*_2_*r*_*g*_). When targeting on the change for dispersion, we let *θ*_1_ = 1 and derive the score function and distribution function for *NB*(*µ*_*g*_, *θ*_2_*r*_*g*_) Since the probability mass function of *NB*(*µ*_*g*_, *θ*_2_*r*_*g*_) is

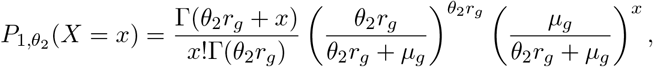

we have

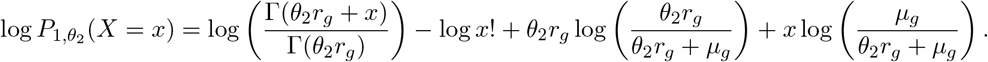

Then the score function at *θ*_2_ = 1 is given by

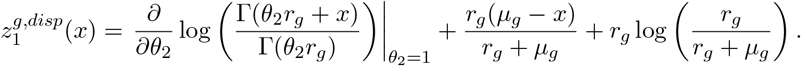

The corresponding Hessian is 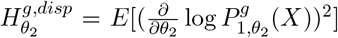, where *X* ∼ *NB*(*µ*_*g*_, *θ*_2_*r*_*g*_). Specifically,

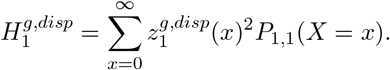

Finally, we also have the CDF of *NB*(*µ*_*g*_, *θ*_2_*r*_*g*_) at *θ*_2_ = 1

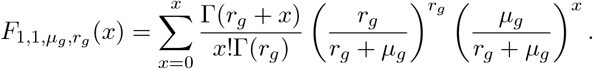

Now, recalling that the probability mass function of *ZINB*(*µ*_*g*_, *θ*_2_*r*_*g*_, *π*_*g*_) is

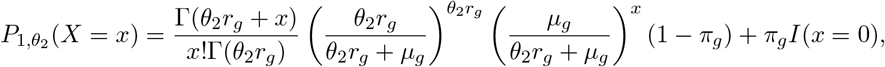

thus we have

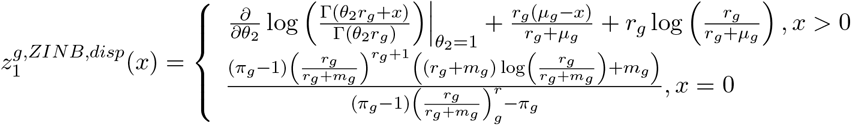

And

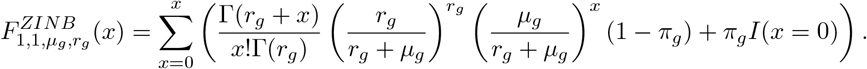

For QRscore-Var, we have 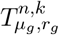 asymptotically follows 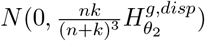.

### A.2 Formula Derivation for Test Statistic in QRscore-Mean

Similarly, we can derive the quantities used to compute the test statistic targeting the mean parameter. Here, we let *θ*_2_ = 1 and derive the score function and distribution function for *NB*(*θ*_1_*µ*_*g*_, *r*_*g*_) Since the probability mass function of *NB*(*θ*_1_*µ*_*g*_, *r*_*g*_) is

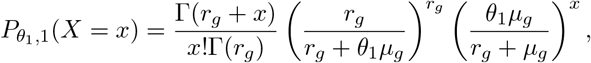

we have

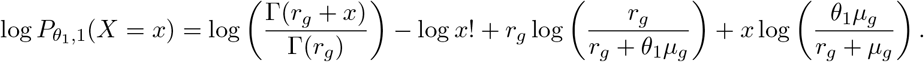

Then the score function at *θ*_1_ = 1 is given by

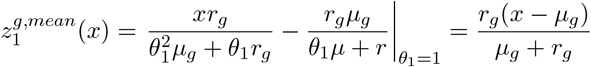

The Hessian for *θ*_1_ = 1 is

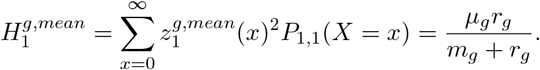

Finally, the CDF of *NB*(*θ*_1_*µ*_*g*_, *r*_*g*_) at *θ*_1_ = 1 is the same as 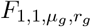.

Now, for *ZINB*(*θ*_1_*µ*_*g*_, *r*_*g*_, *π*_*g*_), thus we have

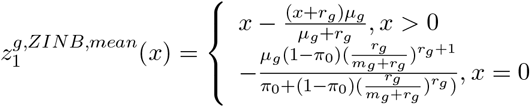

and the CDF 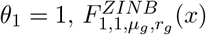 is the same as above.

### A.3 Numerical Considerations and Adjustments for the Test Statistic

Note that the support for the NB distribution is [0, ∞) and the inverse CDF function becomes infinite when 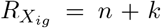.To address this, we made adjustments by replacing *n* + *k* with *n* + *k* + *δ*. Additionally, we centralized the test statistic by subtracting the expected mean under a discrete uniform distribution to eliminate a small bias in the test statistic. The adjusted test statistic, denoted as 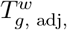, used in practice is given by:

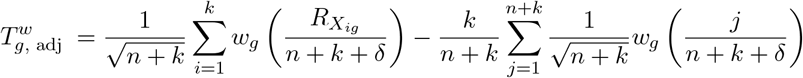

For our analysis, we set *δ* = 1, but *δ* can be any small value greater than 0 to avoid the issue of infinite values from the inverse CDF function. Additionally, we randomly break ties when ranking, ensuring that the rank-based inputs remain unbiased.

## Appendix B *K*-sample test

In Section 4.1.4, we outlined a general approach for *K*-sample testing. Here, we delve into the detailed construction of the test statistic and its inference process specifically for *K* = 3, as applied in our real data analysis. Specifically, for given gene *g*, we set *K* = 3 and designate the samples as *X*_*g*_, *Y*_*g*_, and *Z*_*g*_ with sizes *n*_1_, *n*_2_, and *n*_3_ respectively. The total size is *N* = *n*_1_ + *n*_2_ + *n*_3_. We compute the sample proportions 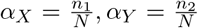,and 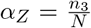.The test statistic is formulated as follows:

**Step 1: Vector Formation** Form a vector of length 3, with its first entry being the sum of the weight function *w* over the ranks of *X*_*g*_, normalized by 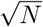. The second and third entries are computed similarly for *Y*_*g*_ and *Z*_*g*_, respectively. The weight function is the same defined in two-sample test, which can be either weights for detecting shifts in mean or weights for detecting shifts in variance.

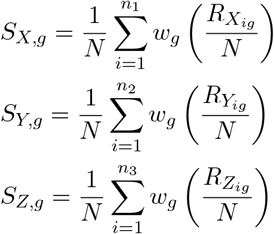

where 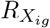 denotes the rank of *X*_*ig*_ after pooling and sorting the values from *X*_*g*_, *Y*_*g*_, and *Z*_*g*_. The ranks 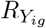 and 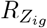 are similarly defined for the values in *Y*_*g*_ and *Z*_*g*_, respectively. Let

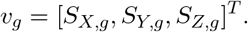

This vector resides in a co-dimension 1 subspace as its entries sum to zero due to the centered score function *w*.

**Step 2: Pre-Multiplication by Projection Matrix** For *K* = 3, a matrix *B* that serves as an orthonormal basis for the subspace can be chosen as follows:

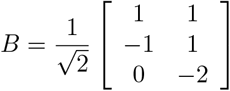

Each column vector in *B* is an orthonormal basis vector of the subspace where the vector from Step 1 resides.

Matrix *A* is a basis for the orthogonal complement of 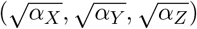.One example for *AA*^*T*^ such that A satisfy such condition is as follows.

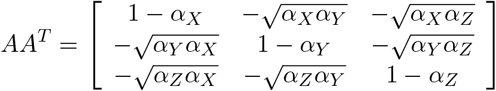

Finally, the diagonal matrix Λ for the *K* = 3 case can be expressed as:

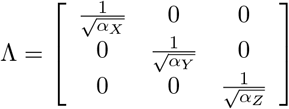

With these matrices, We can proceed with the computation of the test statistic and perform the necessary tests as outlined in the section.

**Step 3: Test statistic construction and inference** We multiply the vector *v*_*g*_ by matrix B and Λ. Let *T*_*g*_ = *B*^*T*^ Λ*v*_*g*_ and 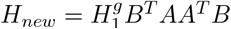.Note that 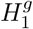 is the same as the Hessian value estimated in the two sample test. According to Theorem 7 in [35], under the null hypothesis that the 3 samples are sampled from the same distribution, we know that the two dimensional random variable *T*_*g*_ asymptotically follows *N* (0, *H*_new_). Therefore, when *n*_1_, *n*_2_, *n*_3_ → ∞,

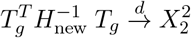

and a chi-square test is applied to the resulting scalar for inference.

## Appendix C

Simulation Details

Our simulation is based on bulk RNA-seq count matrix from whole blood tissue with age groups 40-49 vs. 60-69 from GTEx project. We firstly simulate data with changes in dispersion using negative binomial model. In addition to that, we also designed some simulations that generate different proportion of random zeros or random ourliers to test robustness for all the methods. Furthermore, we simulated few individuals in each age group to check the methods’ sensitivity to sample size.

### C.1 Size factors estimation for normalization

To ensure the integrity of our simulations, we filtered genes with inflated zeros by removing genes with the proportion of zero greater than 20% in any age group. This filtering is only implemented during the setup of the simulation pipeline to prevent the occurrence of inflated zeros if no zeros are deliberately inserted in the simulation. Suppose the expression raw count for each gene *g* and each individual *i* is *M*_*ig*_. We estimated the normalization size factor for each individual i (*s*_*i*_) by taking the median of scaled counts across the genes, which is also applied in DESeq2.

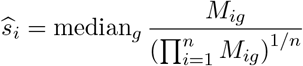

We then normalized the counts by 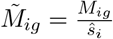.

### C.2 Negative binomial parameters estimation

For each gene, we estimated mean and variance parameters used for simulation from the normalized values for individuals aged 40-49 (*n*_1_ = 133) and 60-69 (*n*_2_ = 249) together with outliers removed. Specifically, let 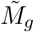 be a vector of normalized expression counts for gene *g*. Let 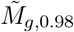 be the normalized counts below the 98th percentile of 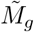.Then we estimate the mean and variance as follows:

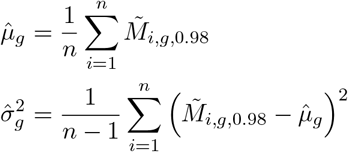

where *n* is the vector length of 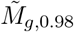,and the dispersion parameter is estimated as 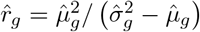.

### C.3 Simulation settings

#### Simulation for dispersion shifts

In the following settings, we simulated N×G count matrices, where *N* = *n*_1_ + *n*_2_ and *G* = 3000. We set the fold change of dispersion parameter (*θ*_2_) to be 1.1, 1.3, …, 2.9 and for each fold change we set 20 random seeds to repeat the experiments.

**Setting 1:** In this simulation, data are generated from a purely negative binomial model with *n*_1_ = 133 individuals in Group 1 (*X*_*ig*_) and *n*_2_ = 249 individuals in Group 2 (*Y*_*jg*_). We randomly select 10% of the genes to be true DDGs, denoted by the set *G*_*D*_. For settings where Group 1 has larger variance than Group 2, we multiply the dispersion parameter by the fold change *θ*_2_. The counts for these genes are simulated as 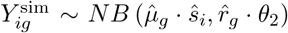 for each *g* ∈ *G*_*D*_. Conversely, for settings where Group 2 has larger variance than Group 1, we divide the dispersion parameter by the fold change, where 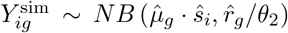.For genes *g* ∈*/G*_*D*_ or individuals in the opposite group (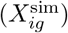,the counts are simulated from 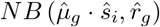 without modification by the fold change. We denote the full simulated matrix that concatenates the two groups as *M* ^sim^.

**Setting 2 - 3:** Simulation from purely negative binomial model for smaller sample sizes. Same as Setting 1, but decrease the number of individuals in groups to *n*_1_ = 50, *n*_2_ = 100 (Setting 2) and *n*_1_ = *n*_2_ = 50 (Setting 3). We random sample *n*_1_ individuals to be in Group 1, and *n*_2_ individuals in Group 2.

**Setting 4 - 6:** Simulation with random outliers. Following the steps in Setting 1, we randomly select 5%, 10%, 20% of the genes to have outliers (*G*_*ol*_). For *g* ∈ *G*_*ol*_, we then select a proportion (*p*_*g*_) of cells with outliers, where *p*_*g*_ ∼ *Unif* (0.25, 0.1). Specifically, the outliers are set to be 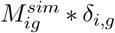 where *δ* ∼ *Unif* (4, 8).

**Setting 7 - 9:** Simulation with random zeros. Following the steps in Setting 1, we randomly select 5%, 10%, 20% of the genes to have inflated zeros (*G*_*zi*_). For *g* ∈ *G*_*zi*_, we select a proportion (*p*_*g*_) of cells to be zeros, where *p*_*g*_ ∼ *Unif* (0.25, 0.1).

**Setting 10:** Simulation with both random outliers and random zeros. Following the steps in Setting 1, we randomly select 10% of the genes to have outliers (*G*_*ol*_) and another 10% of the genes to have inflated zeros (*G*_*zi*_). These selections are made separately, with the possibility that some genes may be included in both groups. The way to generate outliers and zeros are the same as Setting 5 and Setting 7 if genes are in *G*_*ol*_ or *G*_*zi*_.

**Setting 11 - 12:** Simulation with both random outliers and random zeros for smaller sample sizes. Same as Setting 10, but decrease the number of individuals in groups to *n*_1_ = 50, *n*_2_ = 100 (Setting 11) and *n*_1_ = *n*_2_ = 50 (Setting 12).

#### Simulation for mean shifts

In the following settings, we simulated N×G count matrices, where *N* = *n*_1_ + *n*_2_ and *G* = 3000. We set the fold change of mean parameter (*θ*_1_) to be 1.1, 1.2, …, 2 and for each fold change we set 20 random seeds to repeat the experiments.

In Setting 1, we simulate data from a purely negative binomial model with *n*_1_ = 133 individuals in Group 1 and *n*_2_ = 249 individuals in Group 2. We randomly select 10% of the genes to be true DEGs, denoted by the set *G*_*E*_. For each gene *g* ∈ *G*_*E*_, we apply a fold change in dispersion, denoted by *θ*_2_. Specifically, for individuals in Group 1 (*X*_*ig*_), the simulated counts are generated from 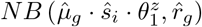,where *z*_*g*_ follows a Rademacher distribution. For genes *g* ∈*/G*_*E*_ or for individuals in Group 2 (*Y*_*jg*_), the simulated counts are generated from 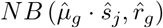.

For other settings, we use the same way to simulate random outliers, zeros or reduced sample sizes. Note that in setting 1 we use the Rademacher distribution to introduce variability in the direction of the mean shifts, rather than fixing the parameter to only increase or decrease. If the mean were increased for all DEGs in Group 2, the size factor estimators would become systematically larger than the true values for all cells in Group 2. This would lead to an artificial mean shift in non-DEGs after normalization, resulting in an inflated number of false discoveries across all models. To avoid this, we randomly select genes to have either increased or decreased variance within one dataset.

### C.4 Evaluation metrics

The definition of FDR and Power are as follows.

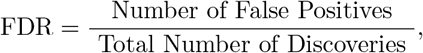

where discoveries are genes with an adjusted *p*-value *<* 0.05.

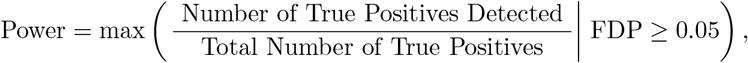

where FDP is the false discovery proportion. In other words, the *p*-value threshold is adaptively chosen for each dataset with the false discovery proportion truly controlled.

For calculating the Area Under the Precision-Recall Curve (AUPRC), Precision and Recall are defined as follows, where the *p*-value threshold is varied continuously from 0 to 1, rather than a fixed threshold:

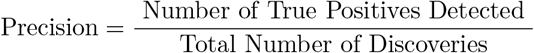

and

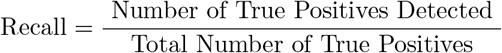

If for each gene, we simulate the same negative binomial distribution for two groups, where all genes follow null hypothesis, the Type I error Rate is computed as follows:

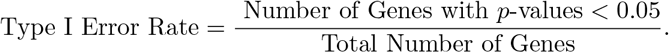

## Appendix D Supplementary Tables

**Table S1:** Average Area Under the Precision-Recall Curve (AUPRC) across all fold changes and all seeds for each simulation setting. The dispersions of Group 2 are greater than those of Group 1 for true DDGs. For each setting, the greatest value is shown in bold. Sample size for Group 1: *n*_1_, sample size for Group 2: *n*_2_, proportion of genes with random outliers: *α*, proportion of genes with random zeros: *β*. Sample sizes *n*_1_ = 113 and *n*_2_ = 249 are chosen from GTEx whole blood tissue data, corresponding to 40-49 and 60-69 age groups respectively. When perturbations are introduced, QRscore outperforms the others, and it ranks second when model assumptions are perfectly met.

| Setting | $n_1$ | $n_2$ | $\alpha$ (outliers) | $\beta$ (zeros) | DiffVar | GAMLSS | Levene | MDSeq | Mood | QRscore-Var |
| --- | --- | --- | --- | --- | --- | --- | --- | --- | --- | --- |
| 1 | 113 | 249 | 0 | 0 | 0.779 | **0.809** | 0.773 | 0.763 | 0.744 | 0.790 |
| 2 | 50 | 100 | 0 | 0 | 0.612 | **0.672** | 0.611 | 0.604 | 0.559 | 0.621 |
| 3 | 50 | 50 | 0 | 0 | 0.555 | **0.603** | 0.541 | 0.516 | 0.492 | 0.576 |
| 4 | 113 | 249 | 0.05 | 0 | 0.774 | 0.768 | 0.769 | 0.688 | 0.742 | **0.785** |
| 5 | 113 | 249 | 0.1 | 0 | 0.767 | 0.734 | 0.762 | 0.633 | 0.740 | **0.781** |
| 6 | 113 | 249 | 0.2 | 0 | 0.756 | 0.675 | 0.753 | 0.547 | 0.738 | **0.777** |
| 7 | 113 | 249 | 0 | 0.05 | 0.752 | 0.749 | 0.760 | 0.763 | 0.739 | **0.782** |
| 8 | 113 | 249 | 0 | 0.1 | 0.725 | 0.698 | 0.743 | 0.764 | 0.733 | **0.778** |
| 9 | 113 | 249 | 0 | 0.2 | 0.664 | 0.605 | 0.708 | 0.760 | 0.719 | **0.762** |
| 10 | 113 | 249 | 0.1 | 0.1 | 0.714 | 0.650 | 0.733 | 0.631 | 0.730 | **0.769** |
| 11 | 50 | 100 | 0.1 | 0.1 | 0.529 | 0.451 | 0.569 | 0.440 | 0.539 | **0.595** |
| 12 | 50 | 50 | 0.1 | 0.1 | 0.472 | 0.389 | 0.500 | 0.363 | 0.472 | **0.542** |

**Table S2:** Average Area Under the Precision-Recall Curve (AUPRC) across all fold changes in mean and all seeds for each simulation setting. For each setting, the highest result is shown in boldface. Sample size for Group 1: *n*_1_, sample size for Group 2: *n*_1_, proportion of random outliers: *α*, proportion of random zeros: *β*.

| Setting | $n_1$ | $n_2$ | $\alpha$ (outliers) | $\beta$ (zeros) | DESeq2 | edgeR | limma | Mann-Whitney | QRscore-Mean |
| --- | --- | --- | --- | --- | --- | --- | --- | --- | --- |
| 1 | 113 | 249 | 0 | 0 | 0.909 | **0.909** | 0.885 | 0.892 | 0.904 |
| 2 | 50 | 100 | 0 | 0 | 0.826 | **0.827** | 0.792 | 0.801 | 0.819 |
| 3 | 50 | 50 | 0 | 0 | 0.786 | **0.787** | 0.750 | 0.760 | 0.775 |
| 4 | 113 | 249 | 0.05 | 0 | 0.906 | **0.906** | 0.873 | 0.890 | 0.903 |
| 5 | 113 | 249 | 0.1 | 0 | 0.903 | **0.903** | 0.858 | 0.888 | 0.902 |
| 6 | 113 | 249 | 0.2 | 0 | 0.899 | 0.899 | 0.833 | 0.888 | **0.901** |
| 7 | 113 | 249 | 0 | 0.05 | 0.898 | 0.898 | 0.883 | 0.891 | **0.902** |
| 8 | 113 | 249 | 0 | 0.1 | 0.887 | 0.887 | 0.882 | 0.890 | **0.900** |
| 9 | 113 | 249 | 0 | 0.2 | 0.868 | 0.868 | 0.879 | 0.889 | **0.897** |
| 10 | 113 | 249 | 0.1 | 0.1 | 0.881 | 0.881 | 0.856 | 0.888 | **0.898** |
| 11 | 50 | 100 | 0.1 | 0.1 | 0.785 | 0.784 | 0.756 | 0.797 | **0.811** |
| 12 | 50 | 50 | 0.1 | 0.1 | 0.739 | 0.738 | 0.710 | 0.752 | **0.763** |

Table S3: Genes with Differential Dispersion Across Multiple Tissues. Table showing genes that exhibit differential dispersion across more than 1 tissue. The table include the gene name, the number of tissues in which differential dispersion is observed, and the specific tissues involved. See attached file: `Sup_table3_multi_tissue_DDGs.xlsx`

Table S4: Coexpressed DDG pairs for each tissue. Supplementary tables showing pairs of DDGs from 20 tissues with significantly enriched GO terms. For each tissue, pairs of DDGs associated with each DDG enriched GO term were analyzed for coexpression using Spearman correlation, with gene pairs having an absolute correlation greater than 0.6 retained. See attached file: `Sup_table4_GO_coexpressed_genes.xlsx`

Table S5: Supplementary tables for 20 tissues with significantly enriched GO terms for DDGs, presenting QRscore-Var and QRscore-Mean results for the top 20 genes associated with the top 10 enriched GO terms. Each table follows the structure of Table 4 and includes data for three age groups in the biological process category as identified by QRscore-Var. See attached file: `Sup_table5_GO_genes.xlsx`

## Appendix E Supplementary Figures

**Figure S1:**
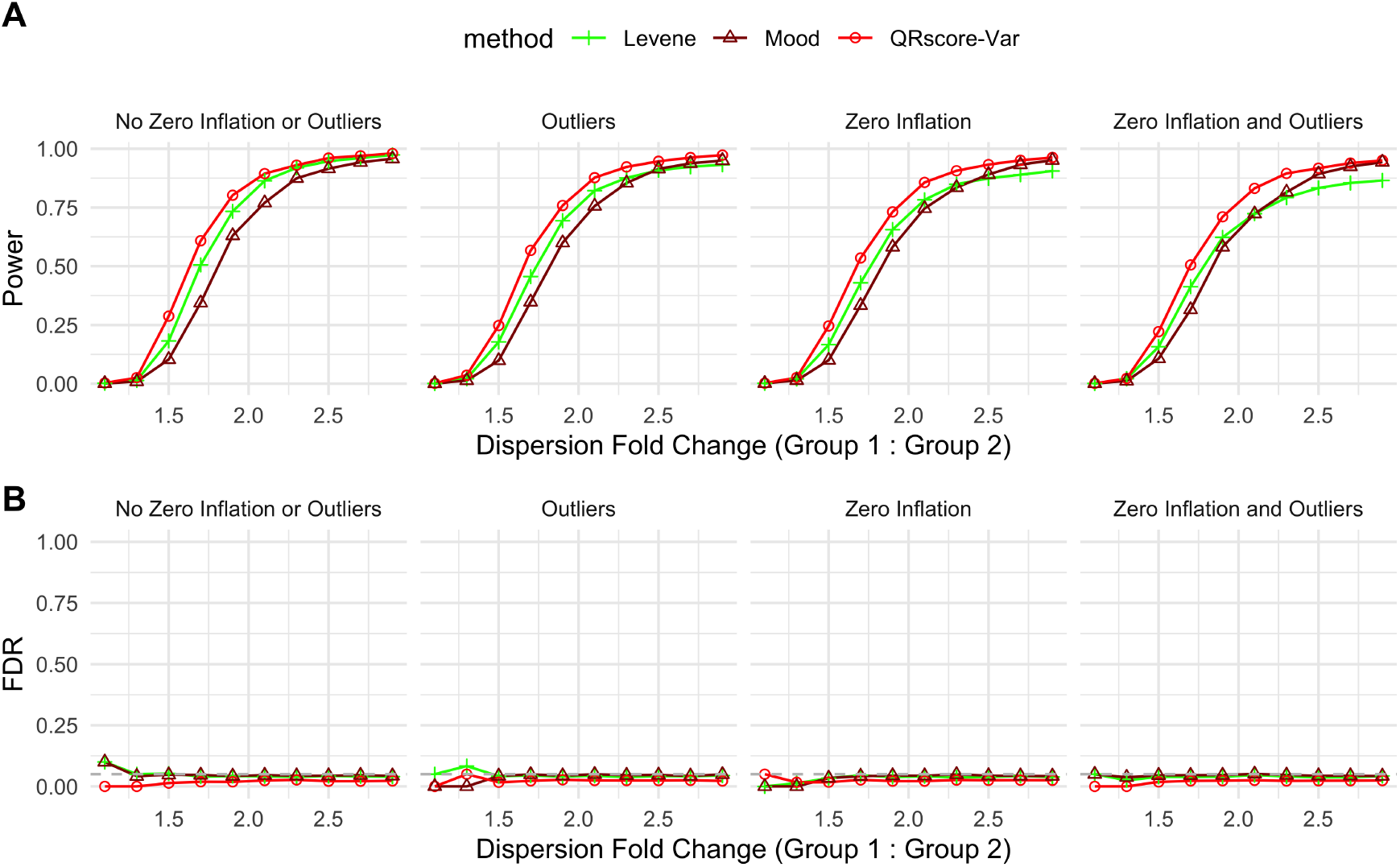
Simulation results for QRscore-Var benchmarked against naive methods for detecting DDGs. The dispersion of group 1 is greater than that of group 2 for true DDGs. (**A**) Line plots of power when true false discovery proportions are less than 0.05 across different dispersion fold changes. (**B**) Line plots of FDR when adjusted *p*-values are less than 0.05 across different dispersion fold changes (Method 4.2). In **A** and **B**, we present the results for sample sizes *n*_1_ = 113 and *n*_2_ = 249. We present four different settings in the plot: one simulated from a pure NB model without random zeros or outliers; one with 10% of genes having outliers; one with 10% of genes having zero inflation; and one with both, where 10% of genes have outliers and 10% have zero inflation.

**Figure S2:**
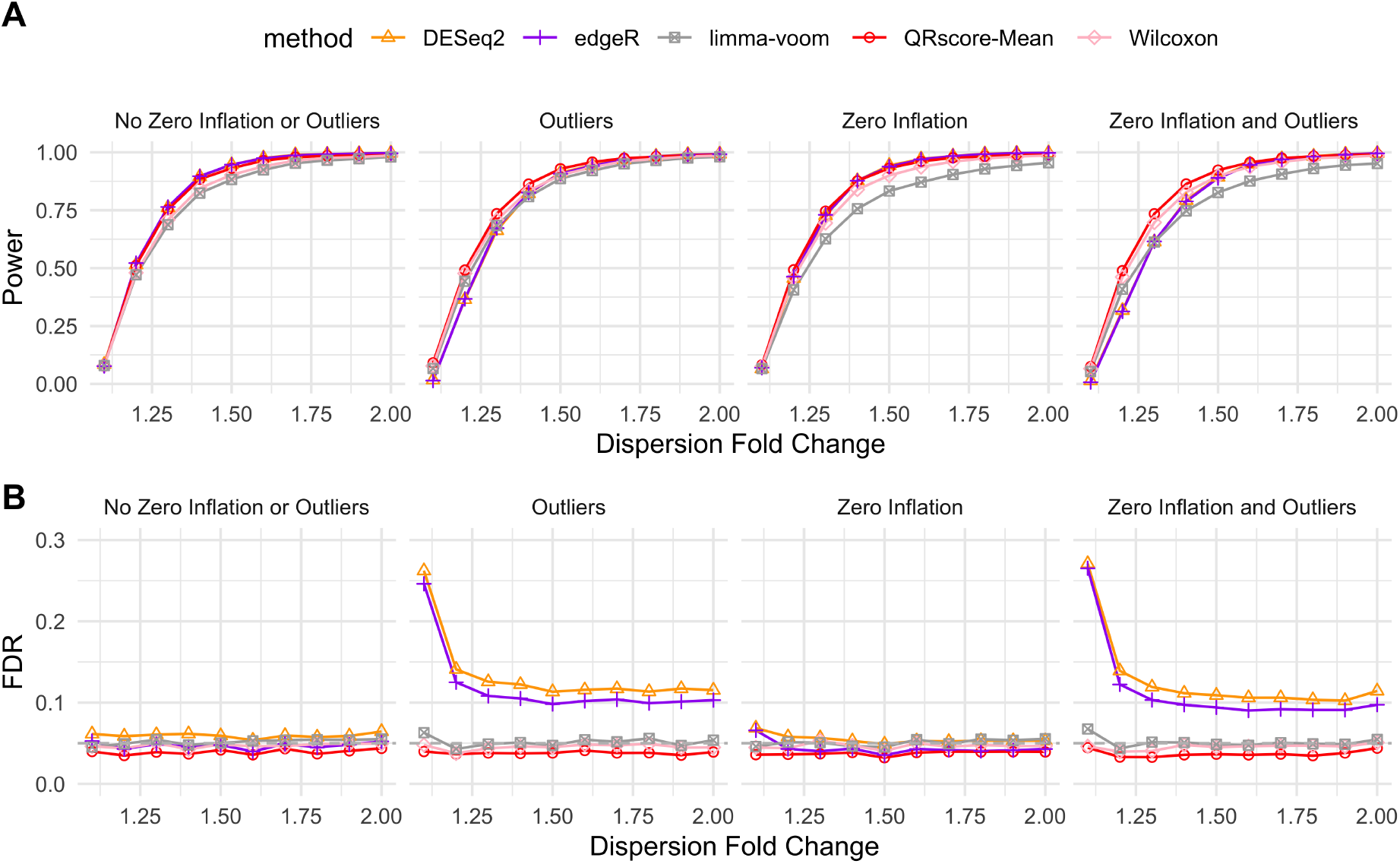
Simulation results for QRscore-Mean. (**A**) Line plots of mean of power when true false discovery proportions (FDP) are less than 0.05 across different mean fold changes. (**B**) Line plots of mean of FDR when adjusted *p*-values are less than 0.05 across different dispersion fold changes. In **A** and **B**, we present the results under Setting 1 (*n*_1_ = 113, *n*_2_ = 249, proportion of random outliers *α* = 0, proportion of random zeros *β* = 0), Setting 5 (*n*_1_ = 113, *n*_2_ = 249, *α* = 10%, *β* = 0), Setting 8 (*n*_1_ = 113, *n*_2_ = 249, *α* = 0, *β* = 10%) and Setting 10 (*n*_1_ = 113, *n*_2_ = 249, *α* = 10%, *β* = 10%).

**Figure S3:**
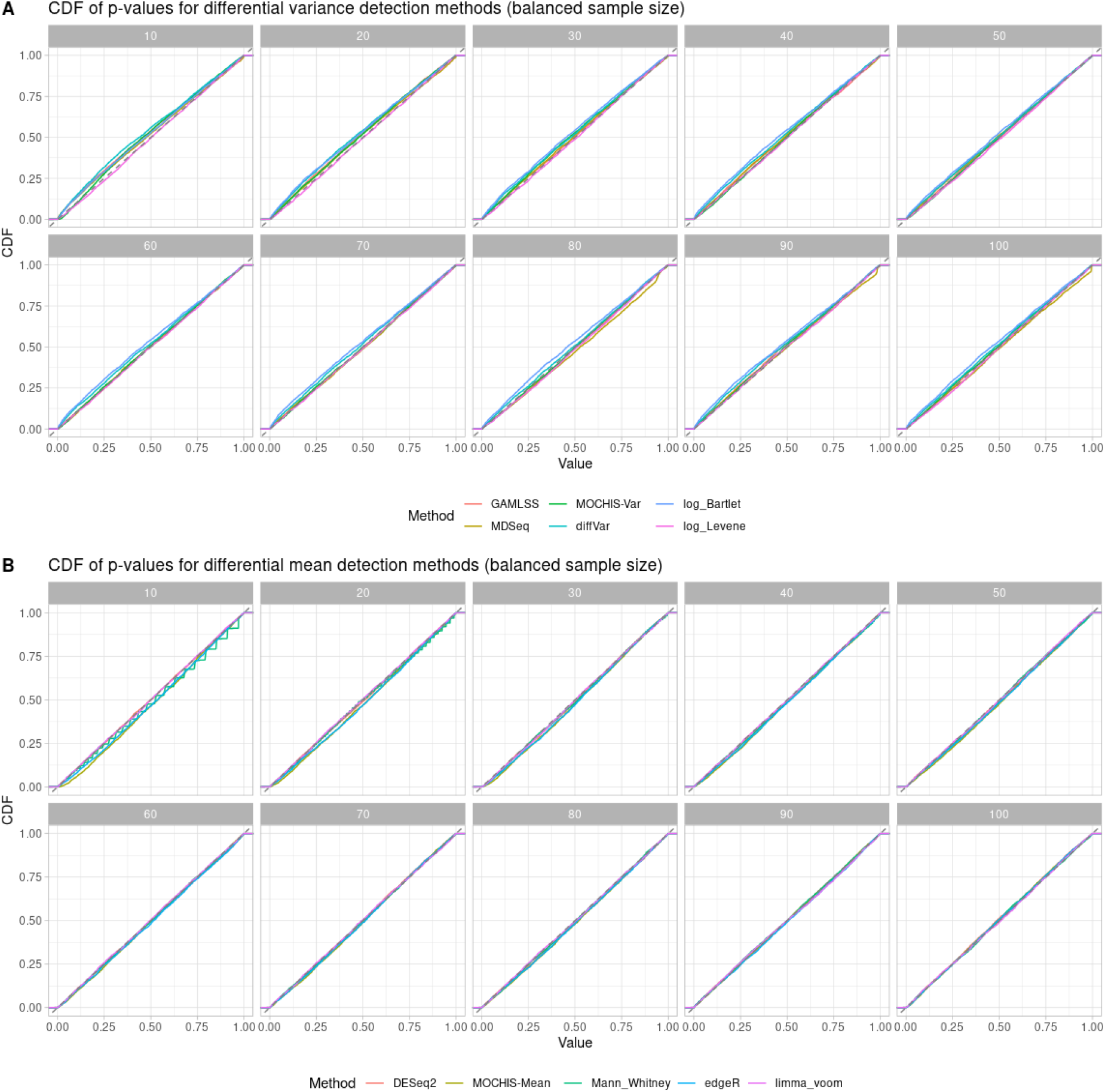
Cumulative distribution function (CDF) of *p*-values from null hypothesis simulations, where data are generated using a pure negative binomial distribution without incorporating any fold change signals. The simulations maintain balanced sample sizes between two groups, varying from 10 to 100.

**Figure S4:**
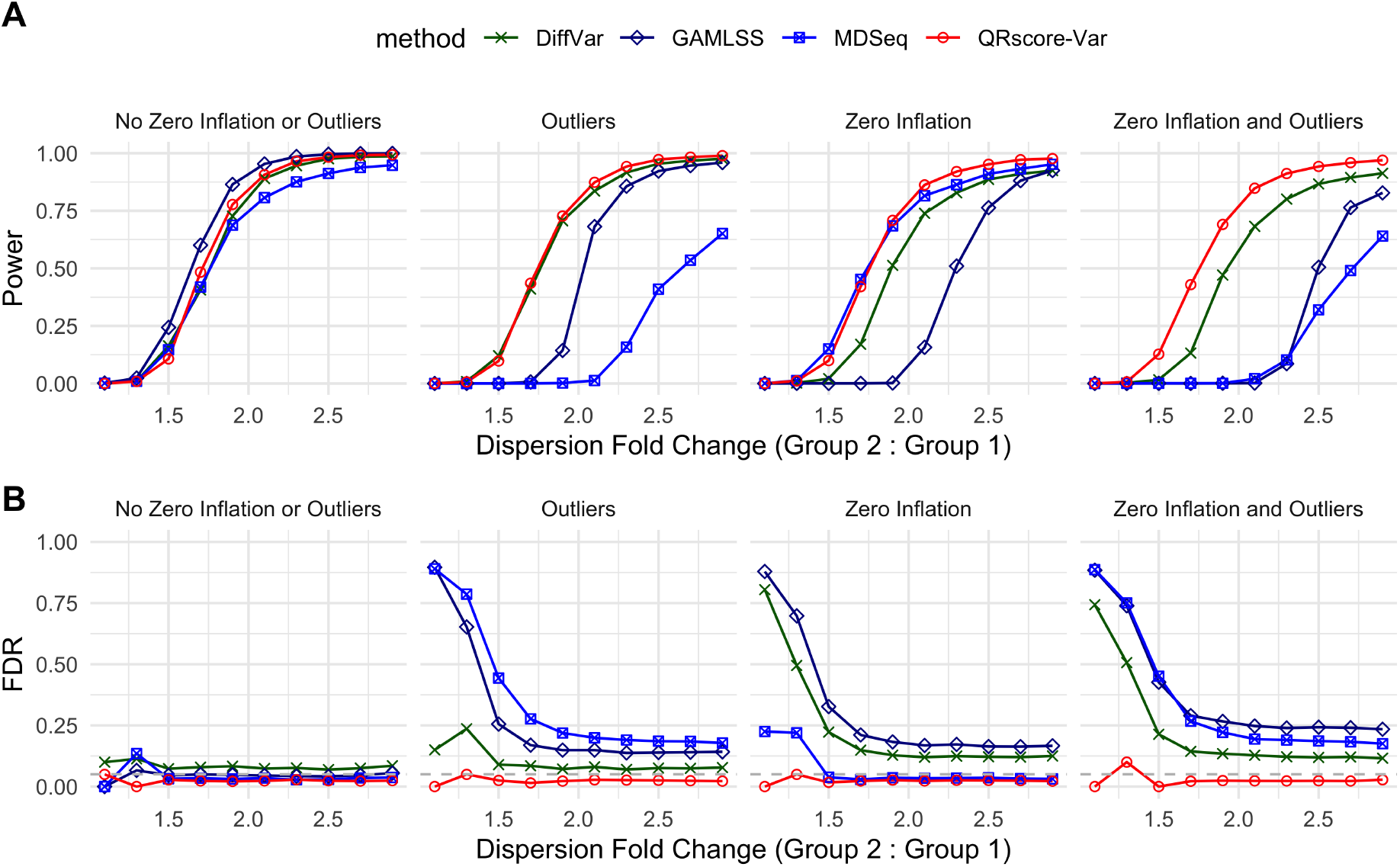
Simulation results for QRscore-Var benchmarked against computational biology methods for detecting DDGs. The dispersion of group 2 is greater than that of group 1 for true DDGs. Line plots of power when true false discovery proportions are less than 0.05 across different dispersion fold changes. (**B**) Line plots of FDR when adjusted *p*-values are less than 0.05 across different dispersion fold changes (Method 4.2). In **A** and **B**, we present the results for sample sizes *n*_1_ = 113 and *n*_2_ = 249. We present four different settings in the plot: one simulated from a pure NB model without random zeros or outliers; one with 10% of genes having outliers; one with 10% of genes having zero inflation; and one with both, where 10% of genes have outliers and 10% have zero inflation.

**Figure S5:**
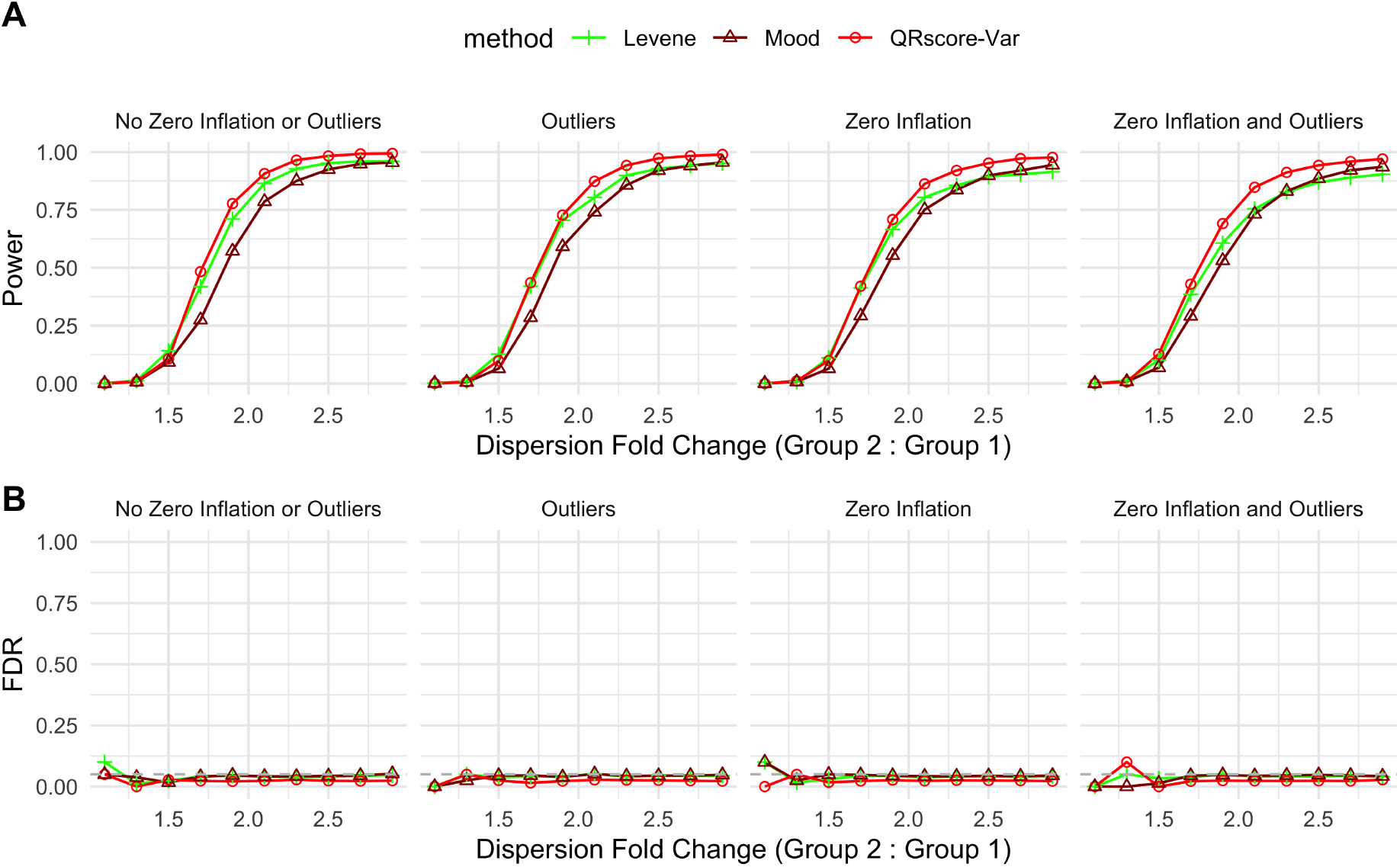
Simulation results for QRscore-Var benchmarked against naive methods for detecting DDGs. The dispersion of group 2 is greater than that of group 1 for true DDGs. (**A**) Line plots of power when true false discovery proportions are less than 0.05 across different dispersion fold changes. (**B**) Line plots of FDR when adjusted *p*-values are less than 0.05 across different dispersion fold changes (Method 4.2). In **A** and **B**, we present the results for sample sizes *n*_1_ = 113 and *n*_2_ = 249. We present four different settings in the plot: one simulated from a pure NB model without random zeros or outliers; one with 10% of genes having outliers; one with 10% of genes having zero inflation; and one with both, where 10% of genes have outliers and 10% have zero inflation.

**Figure S6:**
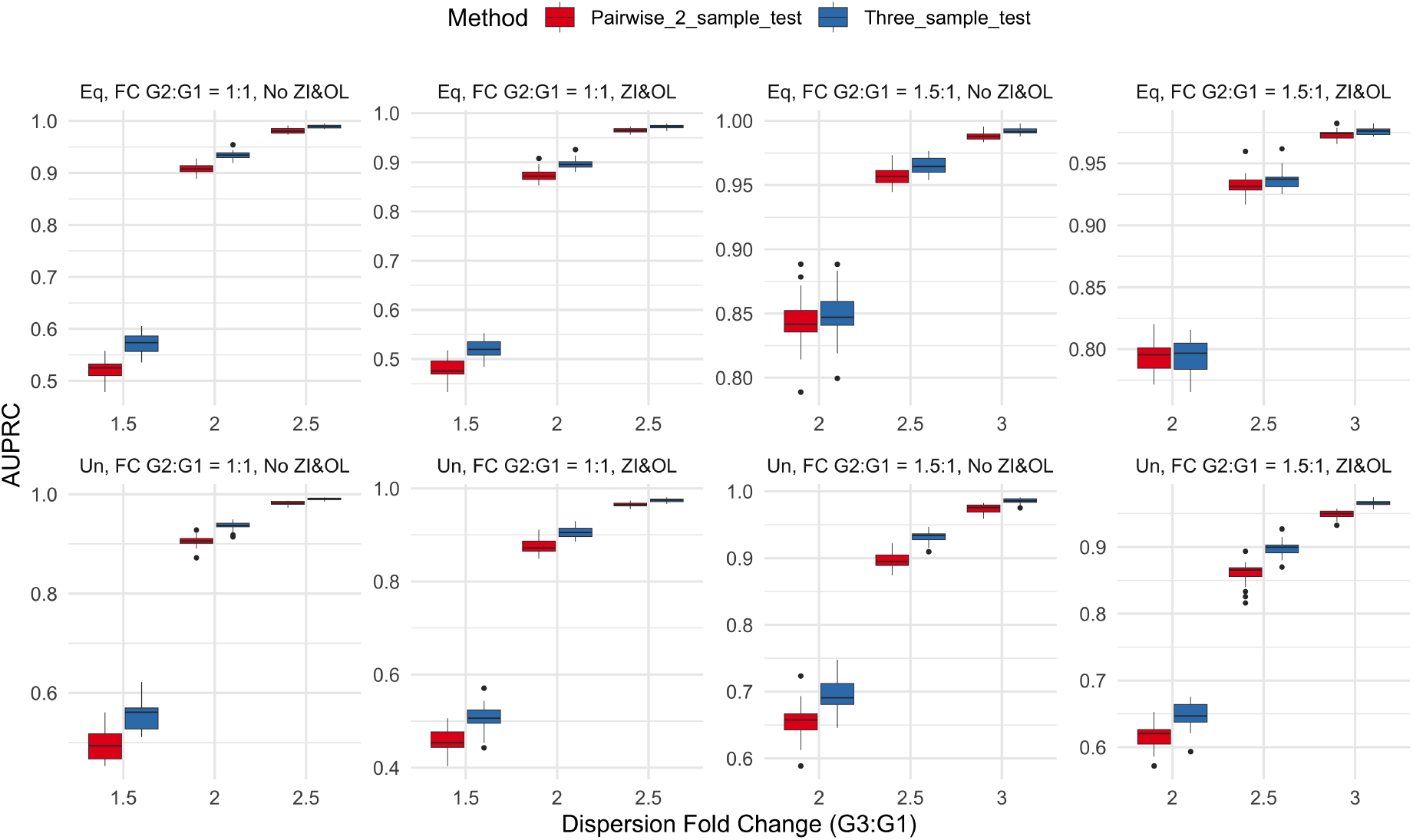
Box plots of AUPRC for pairwise two-sample test and three-sample test using QRscore-Var. The figure presents eight plots depicting different conditions: unequal (Un) sample sizes with *n*_1_ = 50, *n*_2_ = 100, *n*_3_ = 150 and equal (Eq) sample sizes with *n*_1_ = *n*_2_ = *n*_3_ = 100; dispersion fold changes between Group 2 and Group 1 in true DDGs with 1:1 (two samples have the same dispersion, and one sample has different dispersion) and 1:1.5 (all three samples have different dispersions); and the presence of zero inflation and outliers with No ZI&OL (*α* = *β* = 0) and ZI&OL (*α* = *β* = 0.1). Each combination of these conditions is represented, resulting in the eight plots shown.

**Figure S7:**
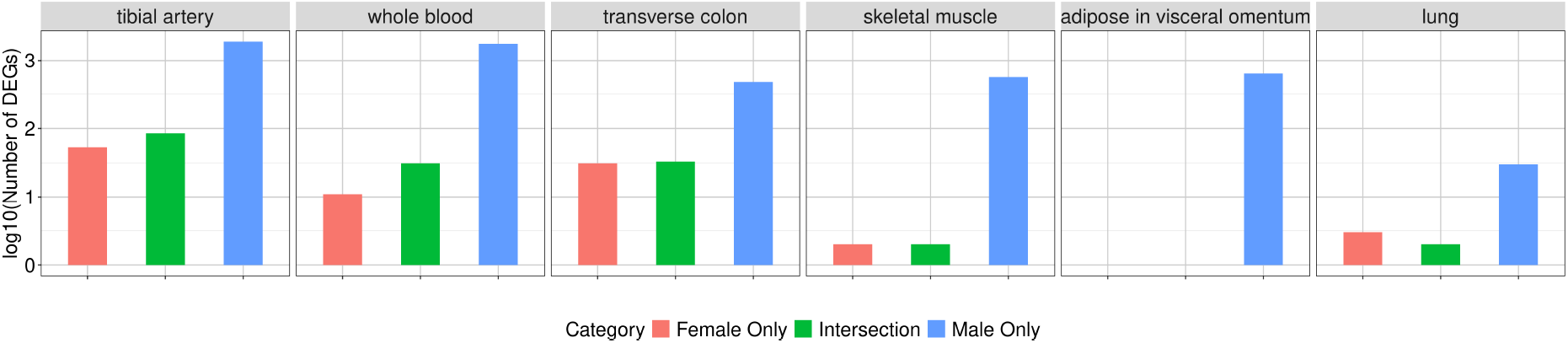
Bar plots display the log counts of DDGs identified in various tissues from male and female individuals, respectively. These results were obtained after stratifying by gender without subsampling the males.

**Figure S8:**
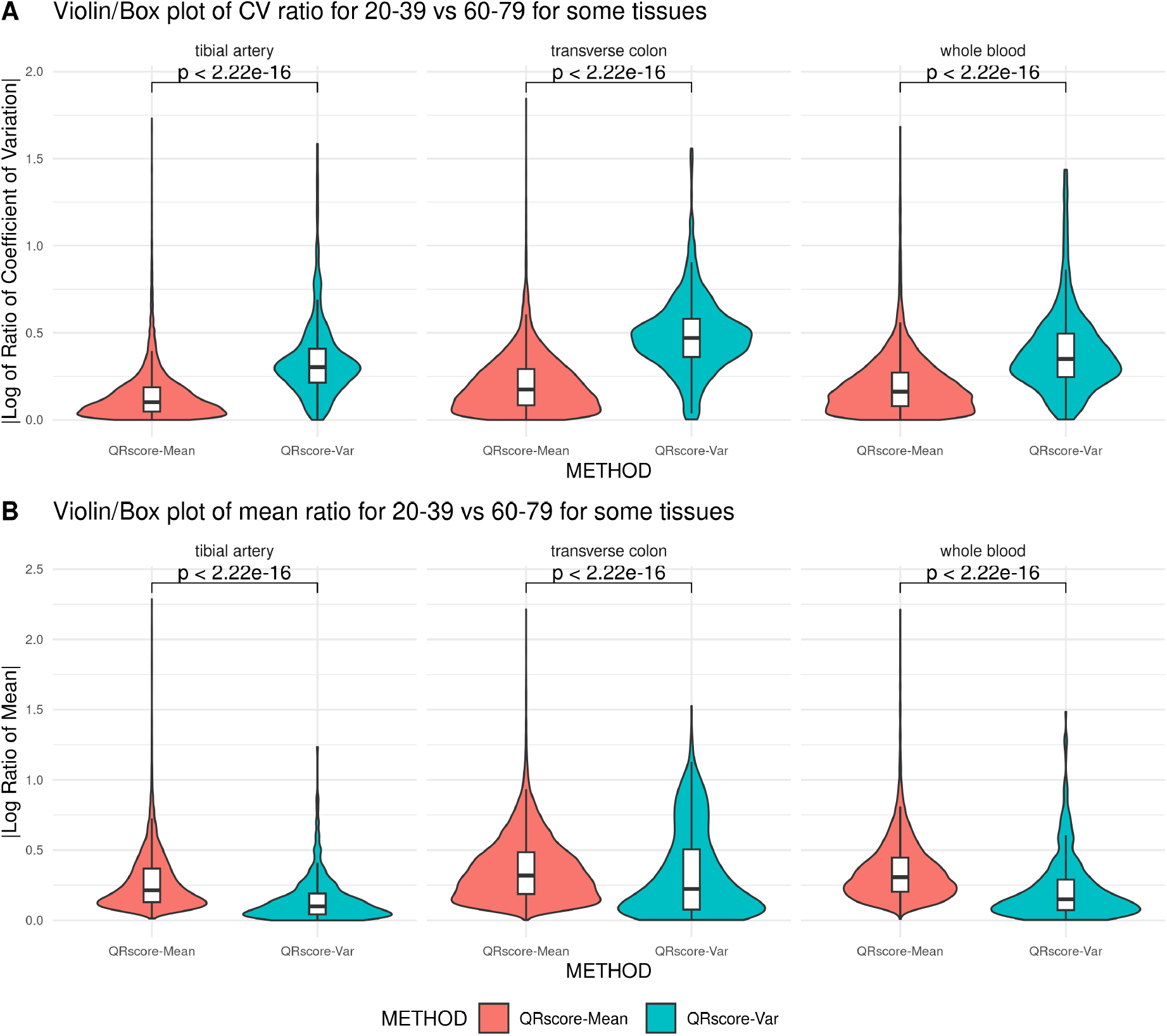
(**A**) Violin and Box plots of log coefficient of variance ratio for genes that only detected by QRscore-Mean and genes that only detected by QRscore-Var from whole blood, transverse colon and tibial artery. (**B**) Violin and Box plots of log mean ratio for genes that only detected by QRscore-Mean and genes that only detected by QRscore-Var from the same tissues in **A**. The DDGs and DEGs are discovered from 3-sample tests and further filtered based on 2-sample tests between 20-39 vs. 60-79 age groups. Extreme outliers (greater than 5 × 3rd quantile) are filtered to get the plots. The *p*-values are obtained from one-sided Wilcoxon test.

**Figure S9:**
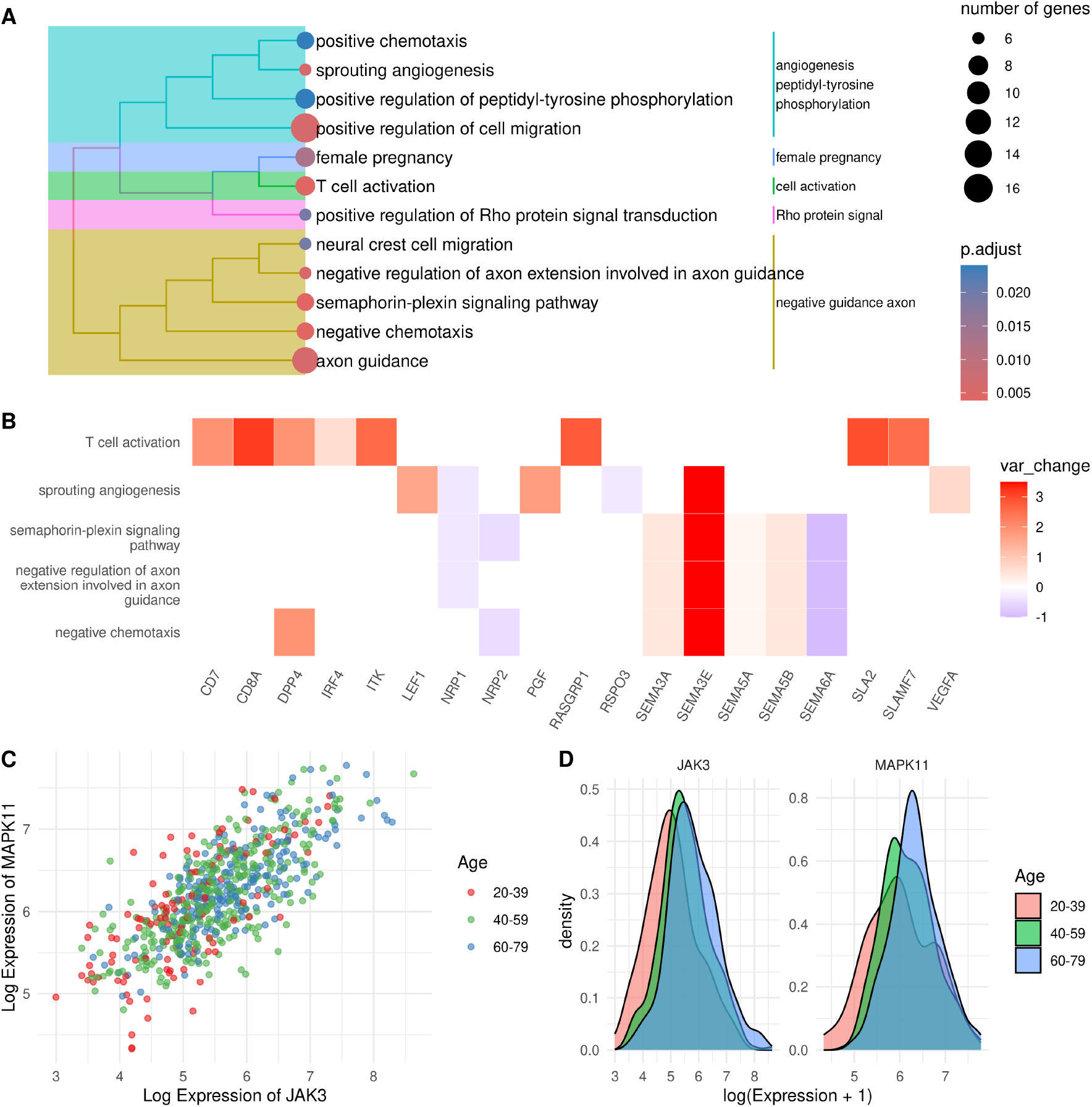
Comparative analysis of GO enrichment in tibial artery tissue based on significant genes identified by QRscore-Var. **(A)** A dot plot displays the top enriched GO terms within the biological process ontology for the top 500 genes significantly detected by QRscore-Var, emphasizing the immune response and T cell activation pathways. **(B)** A heat plot, created using ClusterProfiler, illustrates the fold change in expression variance between the 20-39 and 40-59 age groups for significant genes tied to the top 5 enriched GO terms from the QRscore-Var analysis, highlighting genes involved in immune system processes. **(C)** and **(D)** further explore the expression patterns of CXCL2 and CXCL3, two highly coexpressed genes identified by QRscore-Var as being involved in enriched GO terms, through a dot plot and a density plot, respectively.

**Figure S10:**
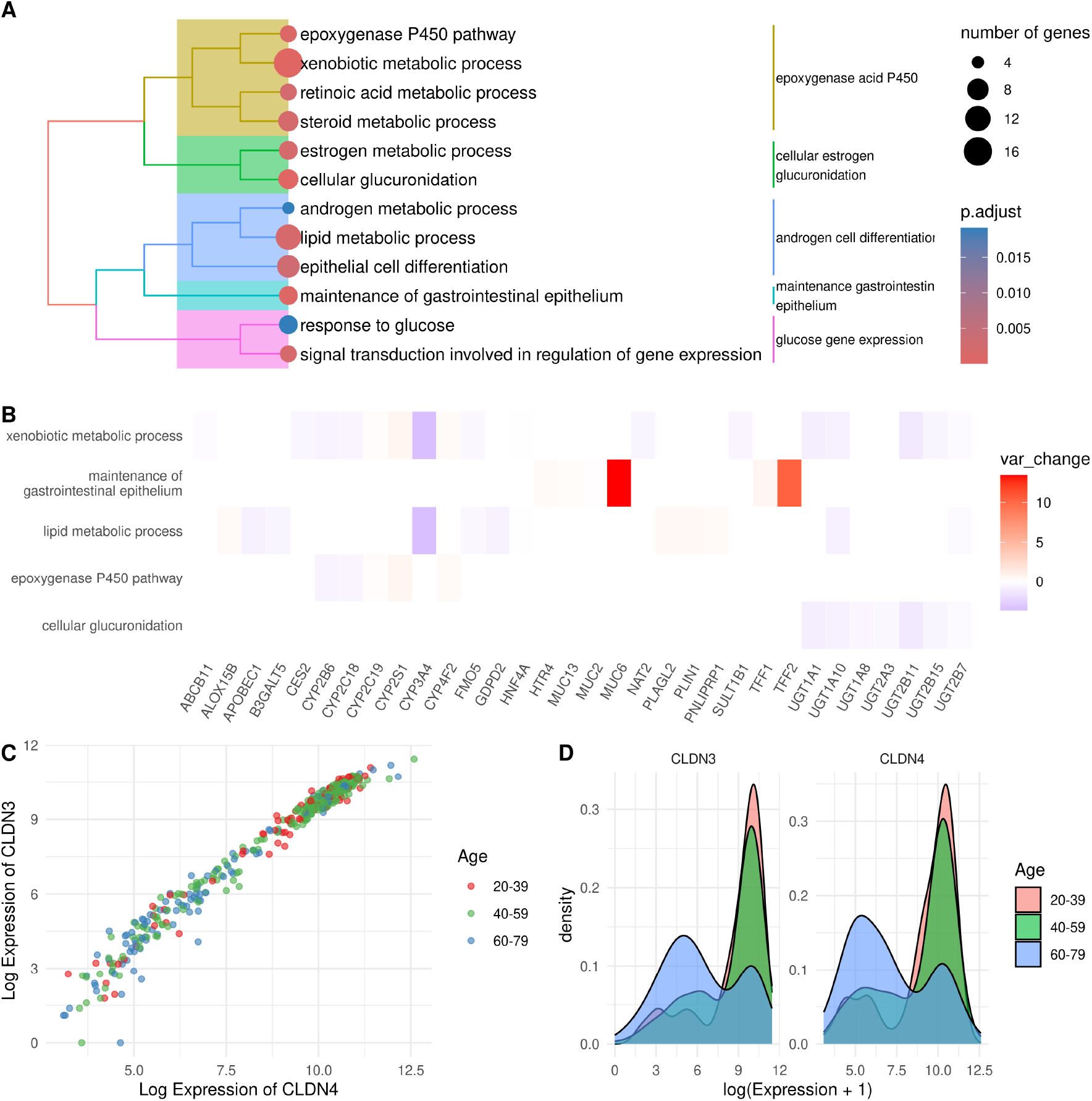
Comparative analysis of GO enrichment in transverse colon tissue based on significant genes identified by QRscore-Var. **(A)** A dot plot displays the top enriched GO terms within the biological process ontology for the top 500 genes significantly detected by QRscore-Var, emphasizing the immune response and T cell activation pathways. **(B)** A heat plot, created using ClusterProfiler, illustrates the fold change in expression variance between the 20-39 and 40-59 age groups for significant genes tied to the top 5 enriched GO terms from the QRscore-Var analysis, highlighting genes involved in immune system processes. **(C)** and **(D)** further explore the expression patterns of CLDN3 and CLDN4, two highly coexpressed genes identified by QRscore-Var as being involved in enriched GO terms, through a dot plot and a density plot.

**Figure S11:**
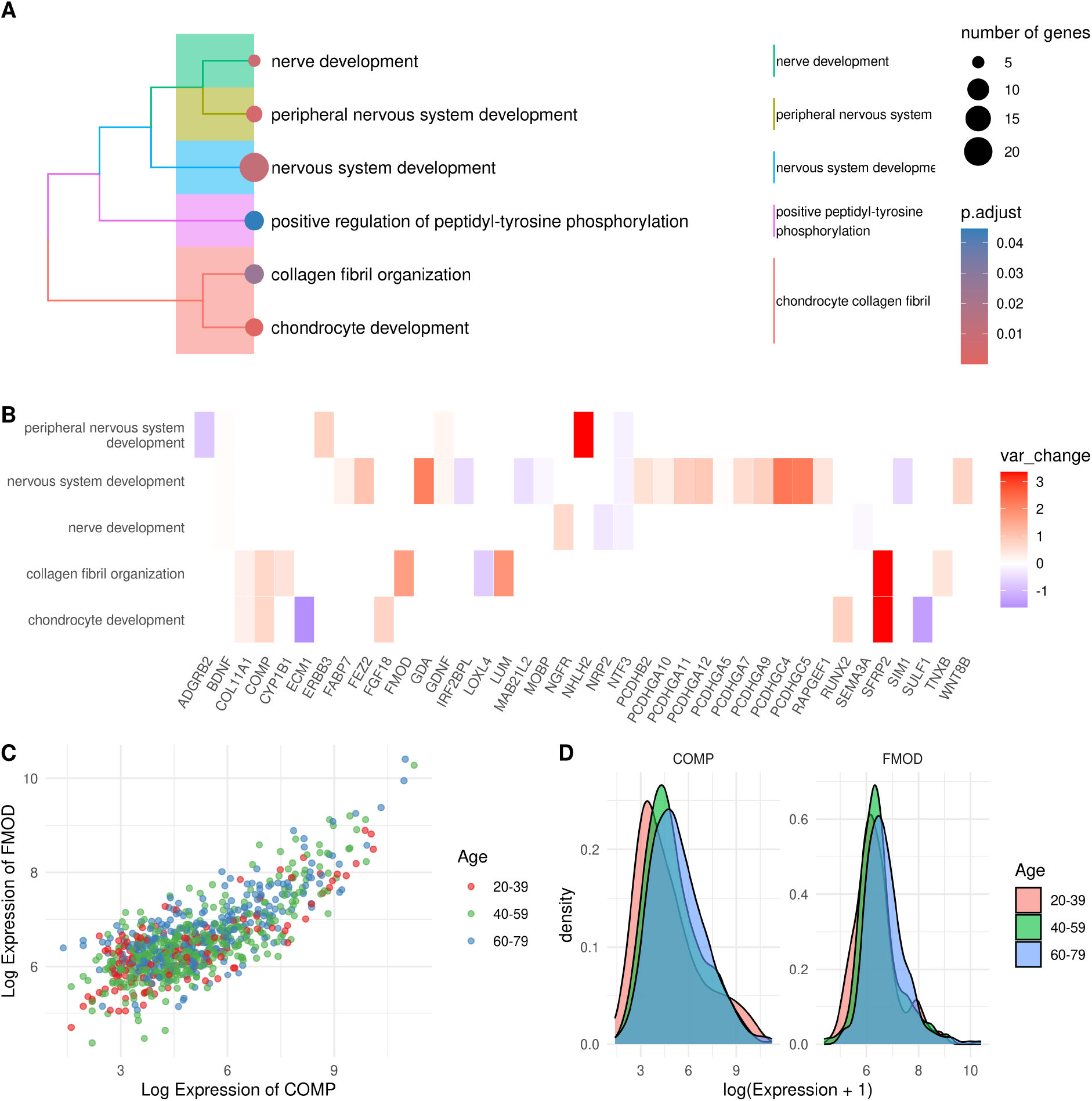
Comparative analysis of GO enrichment in skeletal muscle tissue based on significant genes identified by QRscore-Var. **(A)** A dot plot showcases the top enriched GO terms within the biological process ontology for the top 500 genes significantly detected by QRscore-Var, with a focus on the regulation of vasculature development, angiogenesis, and extracellular matrix organization. These terms highlight the skeletal muscle’s critical roles in vascular health and structural integrity. A heat plot, created using ClusterProfiler, illustrates the fold change in expression variance between age groups for significant genes associated with the top 5 enriched GO terms, emphasizing the dynamic regulation of extracellular structure and angiogenesis with aging. **(C)** and **(D)** examine the expression patterns of COMP and FMOD, two highly coexpressed genes involved in enriched GO terms, through a dot plot and a density plot, respectively.

**Figure S12:**
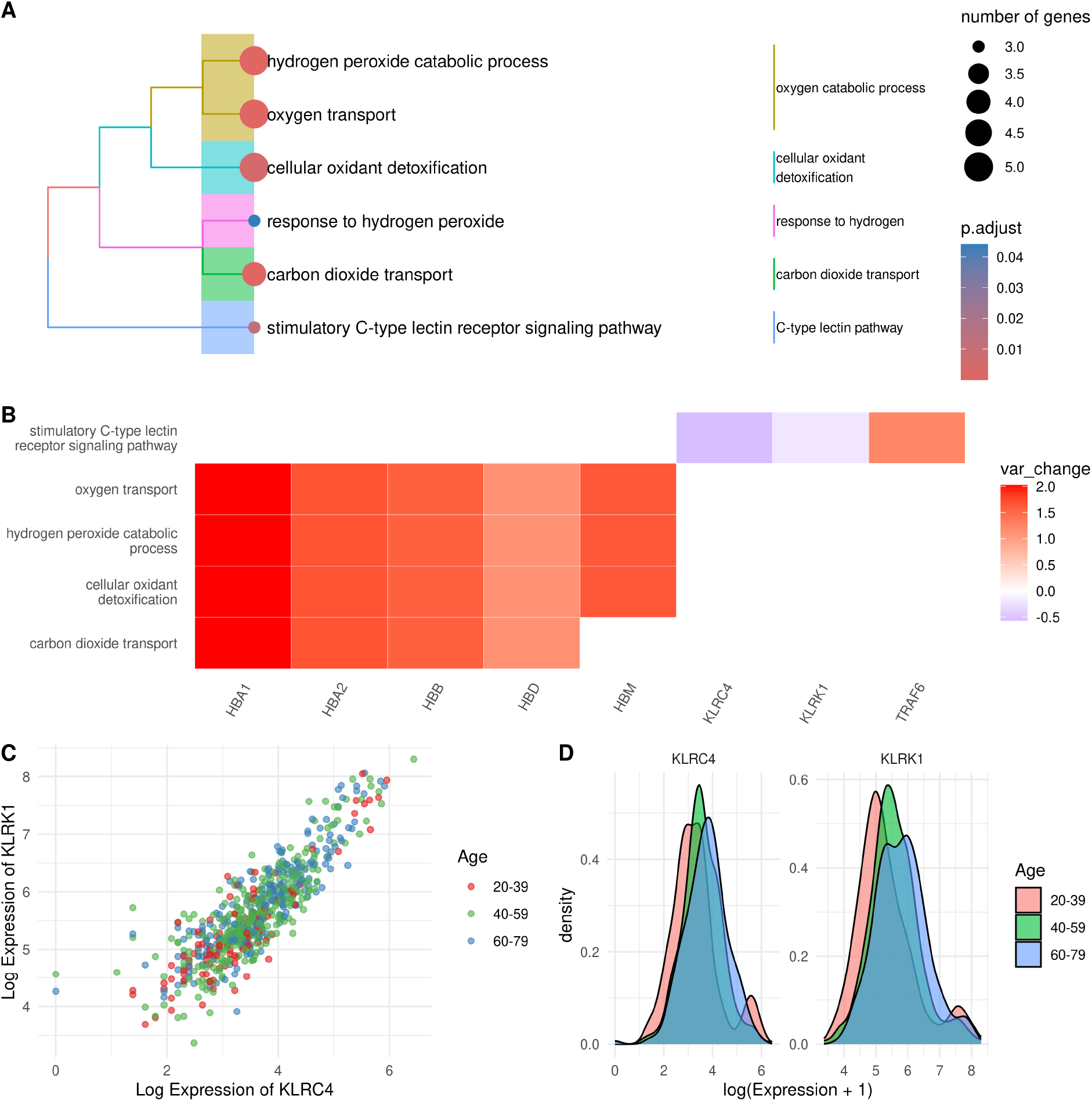
Comparative analysis of GO enrichment in thyroid tissue based on significant genes identified by QRscore-Var. **(A)** A dot plot showcases the top enriched GO terms within the biological process ontology for the top 500 genes significantly detected by QRscore-Var, with a focus on the regulation of vasculature development, angiogenesis, and extracellular matrix organization. These terms highlight the skeletal muscle’s critical roles in vascular health and structural integrity. **(B)** A heat plot, created using ClusterProfiler, illustrates the fold change in expression variance between age groups for significant genes associated with the top 5 enriched GO terms, emphasizing the dynamic regulation of extracellular structure and angiogenesis with aging. **(C)** and **(D)** examine the expression patterns of KLRC4 and KLRC1, two highly coexpressed genes involved in enriched GO terms, through a dot plot and a density plot, respectively.

**Figure S13:**
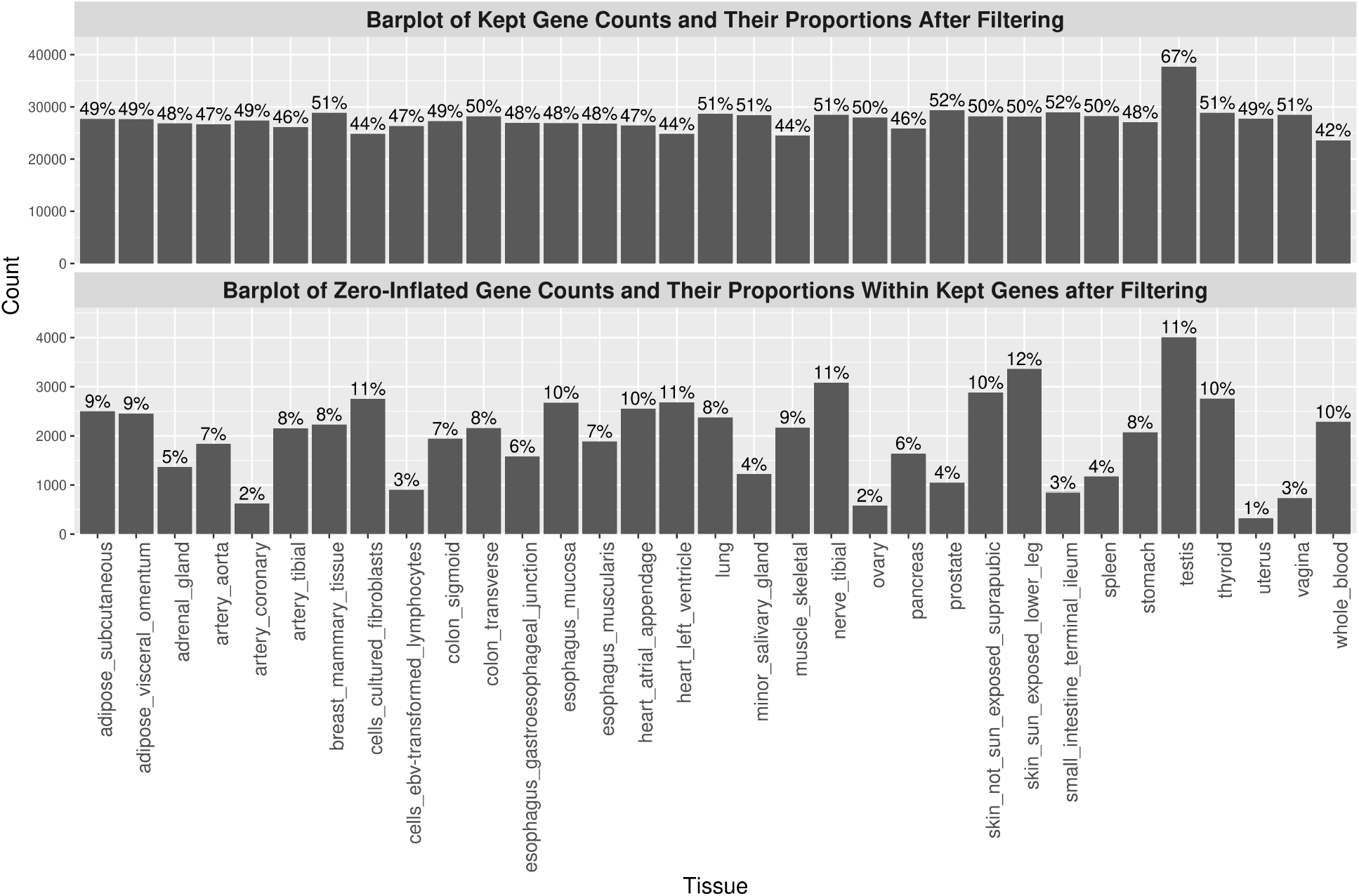
Barplot of number and proportion of genes kept after prefiltering in GTEx dataset and a barplot of zero-inflated gene counts and their proportions within kept genes after filtering. The zero-inflated genes are identified by conducting a likelihood ratio test between the ZINB and NB models fitted for each kept gene, followed by BH adjustment, with an adjusted *p*-value *<* 0.05. Thousands of kept genes in each tissue have inflated zeros, so considering for zero-inflation is still necessary after filtering.

